# Please Mind the Gap: Indel-Aware Parsimony for Fast and Accurate Ancestral Sequence Reconstruction and Multiple Sequence Alignment including Long Indels

**DOI:** 10.1101/2024.03.27.586611

**Authors:** Clara Iglhaut, Jūlija Pečerska, Manuel Gil, Maria Anisimova

## Abstract

Despite having important biological implications, insertion and deletion (indel) events are often disregarded or mishandled during phylogenetic inference. In multiple sequence alignment, indels are represented as gaps and are estimated without considering the distinct evolutionary history of insertions and deletions. Consequently, indels are usually excluded from subsequent inference steps, such as ancestral sequence reconstruction and phylogenetic tree search.

Here, we introduce indel-aware parsimony (indelMaP), a novel way to treat gaps under the parsimony criterion by considering insertions and deletions as separate evolutionary events and accounting for long indels. By identifying the precise location of an evolutionary event on the tree, we can separate overlapping indel events and use affine gap penalties for long indel modelling. Our indel-aware approach harnesses the phylogenetic signal from indels, including them into all inference stages.

Validation and comparison to state-of-the-art inference tools on simulated data show that indelMaP is most suitable for densely sampled datasets with closely to moderately related sequences, where it can reach alignment quality comparable to probabilistic methods and accurately infer ancestral sequences, including indel patterns. Due to its remarkable speed, our method is well-suited for epidemiological datasets, eliminating the need for downsampling and enabling the exploitation of the additional information provided by dense taxonomic sampling. Moreover, indelMaP offers new insights into the indel patterns of biologically significant sequences and advances our understanding of genetic variability by considering gaps as crucial evolutionary signals rather than mere artefacts.

## 1 Introduction

A central aspect of phylogenetics is identifying and understanding the mechanisms driving genetic evolution. While the focus has historically been on substitution events, the second most important source of genetic variation, insertion and deletion (indel) events, are often ignored. Indels are crucial in shaping genomic diversity in various organisms and are closely linked to critical factors such as immune escape, drug resistance, and adaptive selection. For instance, in pathogens like HIV-1 and the malaria parasite *P. falciparum*, indels contribute to the very high genomic diversity and are associated with immune evasion and drug resistance (Wood et al., 2009; Menendez-Arias et al., 2006; Miles et al., 2016). In the ongoing context of the SARS-CoV-2 pandemic, the genomic variants of concern notably exhibit heightened indel rates, suggesting adaptive selection (Alisoltani et al., 2022; Rao et al., 2021). In tumour genomes, indels can serve as informative biomarkers with implications for prognostics and risk stratification (Wu et al., 2019). Furthermore, indels causing variation in protein length, often overlooked in protein evolution studies, hold evolutionary significance and offer opportunities for protein engineering (Savino et al., 2022). Indels are generally known to have a limited impact on the structurally conserved regions of proteins. However, they can exert influence on the variable regions, potentially affecting protein-protein interactions(Studer et al., 2013).

Despite the significance of indels, most phylogenetic inference methods do not systematically reconstruct indel events. During multiple sequence alignment (MSA), indels are represented as gaps and are usually placed according to an optimality score based on the substitution model without considering that gaps represent two distinct evolutionary events, i.e. a gap can be an insertion or a deletion. Commonly used alignment tools, such as MAFFT (Katoh, 2002), hence overlook the nuanced evolutionary context of indels, leading to potential inaccuracies in downstream analyses (Löytynoja and Goldman, 2008; Wong et al., 2008). Consequently, indels are frequently excluded from subsequent inference steps by treating gaps as missing characters during ancestral sequence reconstruction (ASR) or masking gap-rich regions during phylogenetic tree search.

Nevertheless, promoted by the growing recognition of the importance of indels, there has been an increase in attention and exploration of their role in evolutionary analysis, which has subsequently led to the development of indel-aware methods striving to leverage their significant phylogenetic signal (Birth et al., 2022; Dessimoz and Gil, 2010). For example, the MSA tool PRANK (Löytynoja, 2014) uses an outgroup sequence to distinguish between insertion and deletion events during progressive alignment. Additionally, ASR methods such as FastML (Ashkenazy et al., 2012) and GRASP (for amino acid sequences) (Foley et al., 2019) now possess the capability to reconstruct ancestral sequences, including indel events. Notably, ProPIP (Maiolo et al., 2021) and ARPIP (Jowkar et al., 2022), an MSA and ASR method, respectively, adopt an explicit indel model for single residue indel events, utilising the Poisson Indel Process (PIP) (Bouchard-Côté and Jordan, 2013). PIP, a modification of the TKF91 model (Thorne et al., 1991), has enabled the computation of marginal likelihood for an evolutionary model incorporating indel events in linear time with respect to the number of sequences. Furthermore, Historian (Holmes, 2017), a newer implementation of ProtPal Westesson et al. (2012), can be used for indel-aware MSA and ASR estimation. Historian employs a time-dependent evolutionary model and progressively builds an ancestral sequence profile including suboptimal alignments.

In this context, we propose indelMaP, an indel-aware parsimony approach that includes insertions and deletions as distinct events in all inference steps (available at https://github.com/acg-team/indelMaP). This approach allows for a more comprehensive understanding of evolutionary relationships and ancestral sequence reconstruction, in contrast to conventional parsimony methods that treat the gap character as an ambiguous or additional state. Although maximum likelihood and Bayesian methods are commonly preferred for phylogenetic inferences, now, with increasingly growing dataset sizes, parsimony analysis is becoming more popular again, especially for the analysis of very large datasets (Thornlow et al., 2021; Hausdorf, 2022; Bezemer et al., 2022; Torres et al., 2022). Besides being used as a principle in standalone methods (Turakhia et al., 2021; Ye et al., 2022), parsimony can also be utilised as a filter to speed up more complex probabilistic methods (Guindon et al., 2010; De Maio et al., 2023). By including insertion and deletions in the estimation, indel-aware parsimony could be used as a computational approximation for methods that explicitly model indels, which come with a significant computational burden.

## 2 A New Approach: Treating Gaps as Distinct Insertion and Deletion Events under Parsimony

Indel-aware parsimony is an extension of Fitch’s algorithm (Fitch, 1971), which only applies to substitution events. The possible ancestral reconstructions for each site are enumerated in so-called parsimony sets. A union between two parsimony sets that do not share common characters represents a substitution event, while a non-empty intersection between sets represents no evolutionary change. The minimum cost of evolutionary change per site is stored in the parsimony score. To treat indel events as two distinct evolutionary events, we infer the insertion point for each residue based on Dollo’s Law (Farris, 1977), i.e. once a character is deleted, it cannot reappear. The Dollo parsimony insertion point is defined as the most recent point on the tree where the character could have been inserted. Gaps in the subtree defined by the Dollo parsimony insertion point are treated as deletions. In contrast, gaps outside the subtree signify an absence of the character in this part of the tree and are thus merely placeholders. The possible internal reconstructions vary for the two events. If a character is inserted along the current branch, the ancestral parsimony set only holds the gap character, symbolised by the character ‘*’, since inserted characters do not have homologous ancestral characters. On the contrary, if a deletion occurs, the possible internal reconstructions are only residues.

Consequently, we can locate and identify the evolutionary event on the phylogenetic tree, which enables the separation of overlapping indel events and makes it possible to use an affine gap scoring scheme to model long indels.

### 2.1 Inferring indel events and their evolutionary homology

For a fixed MSA and phylogenetic tree, each site’s Dollo parsimony insertion point is located on the branch leading to the most recent common ancestor of all leaves containing a residue at the site (Fig. 1A). A post-order tree traversal calculates each site’s parsimony score and parsimony sets. A subsequent pre-order tree traversal is needed to find the most parsimonious reconstruction for each internal site. Compared to a conventional parsimony method that includes gaps as an additional state (Fig. 1B), indel-aware parsimony successfully identifies and scores each of the evolutionary events represented by the alignment column. A detailed algorithm description can be found in Section 5.1.

**Figure 1.**
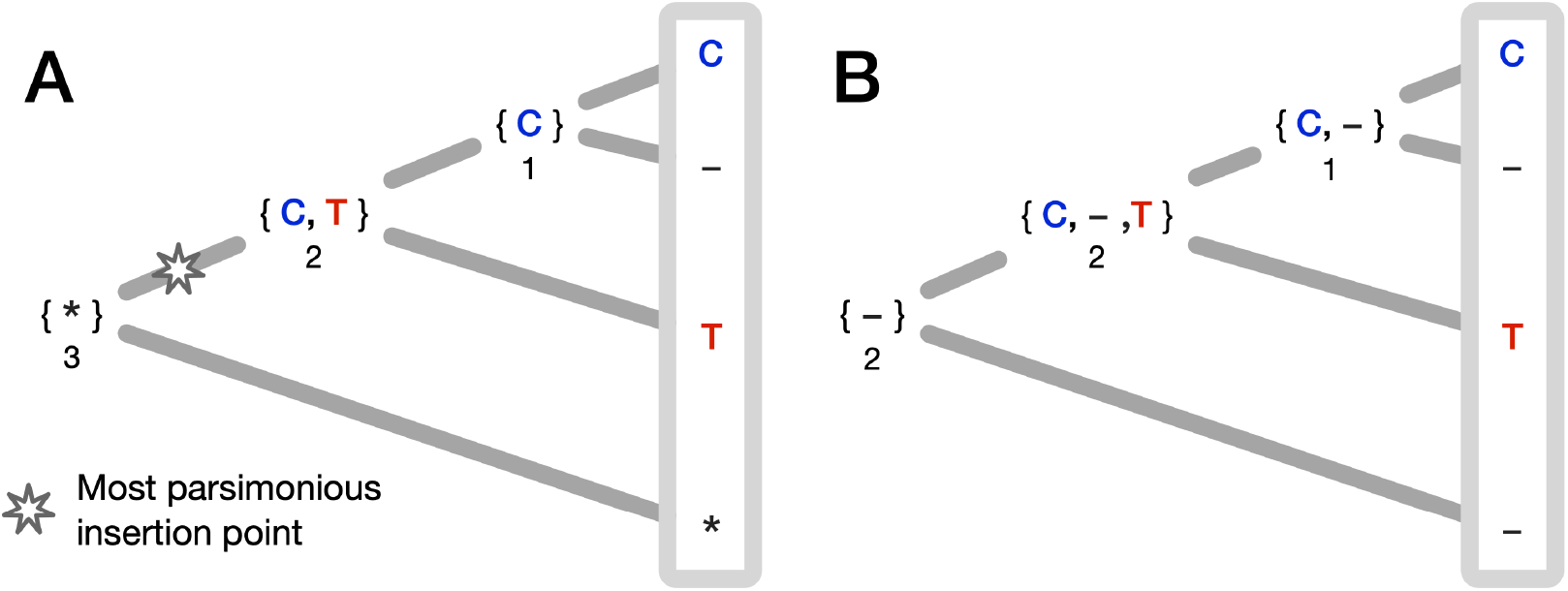
Indel-aware parsimony (A) and parsimony with gaps as an additional state (B) for a fixed alignment column and tree. The parsimony score that memorises the amount of evolutionary change is depicted under each internal node. (A) After identifying the Dollo parsimony insertion point, indel-aware parsimony can separate the three evolutionary events represented by the alignment column. Following the pre-order tree traversal, the site at the root node can unambiguously be reconstructed by the gap character ‘*’, which acts as a placeholder for an insertion event further down the tree. To reconstruct the character at the internal child of the root node, we have to make a random choice between characters C and T. Depending on the choice, either a substitution from C to T or from T to C is reconstructed. The reconstruction for the last node is C, which is deleted in one of the leaf nodes. The total score for the alignment column is three for three events. (B) The conventional method comes to a different conclusion regarding the evolutionary history of the site. Each internal state can unambiguously be reconstructed by a gap character, reconstructing two independent insertions at the same site. Since an alignment column represents homology, and two insertions cannot be homologous to each other, the ancestral reconstruction and the associated parsimony score are not evolutionary meaningful.

Separating insertion and deletions during alignment and inferring a Dollo parsimony insertion point is more challenging. Conventional alignment methods repeatedly penalise insertion sites and fail to place independent insertion events into their individual column since progressive alignment methods cannot differentiate if the gap placement represents an insertion or a deletion in the current node (Löytynoja and Goldman, 2005). We adopted a flagging method previously introduced in (Löytynoja, 2014) to handle indels as two separate events. The flagging method and internal reconstructions used for alignment are illustrated in Fig. 2. Sites aligned with a gap column are marked as a deletion (unfilled orange rhombus in Fig. 2) and reconstructed accordingly. In the subsequent alignment step, the column can be matched with a column containing a residue or a gap column without penalisation. If the former is the optimal decision, we remove the mark for deletion. Otherwise, we mark the site as an insertion site (filled orange rhombus in Fig. 2). Sites marked as insertions are always aligned with a gap column and skipped by the algorithm, i.e., they do not constitute an open gap for the affine gap penalty in later alignment steps. Since we reconstructed the site as a deletion in the previous step, which is now an insertion which happened beneath that node, we have to change the internal reconstruction to a gap character. Note that this is only important if we want to reconstruct ancestral states. A complete description of the algorithm can be found in Section 5.2.

**Figure 2.**
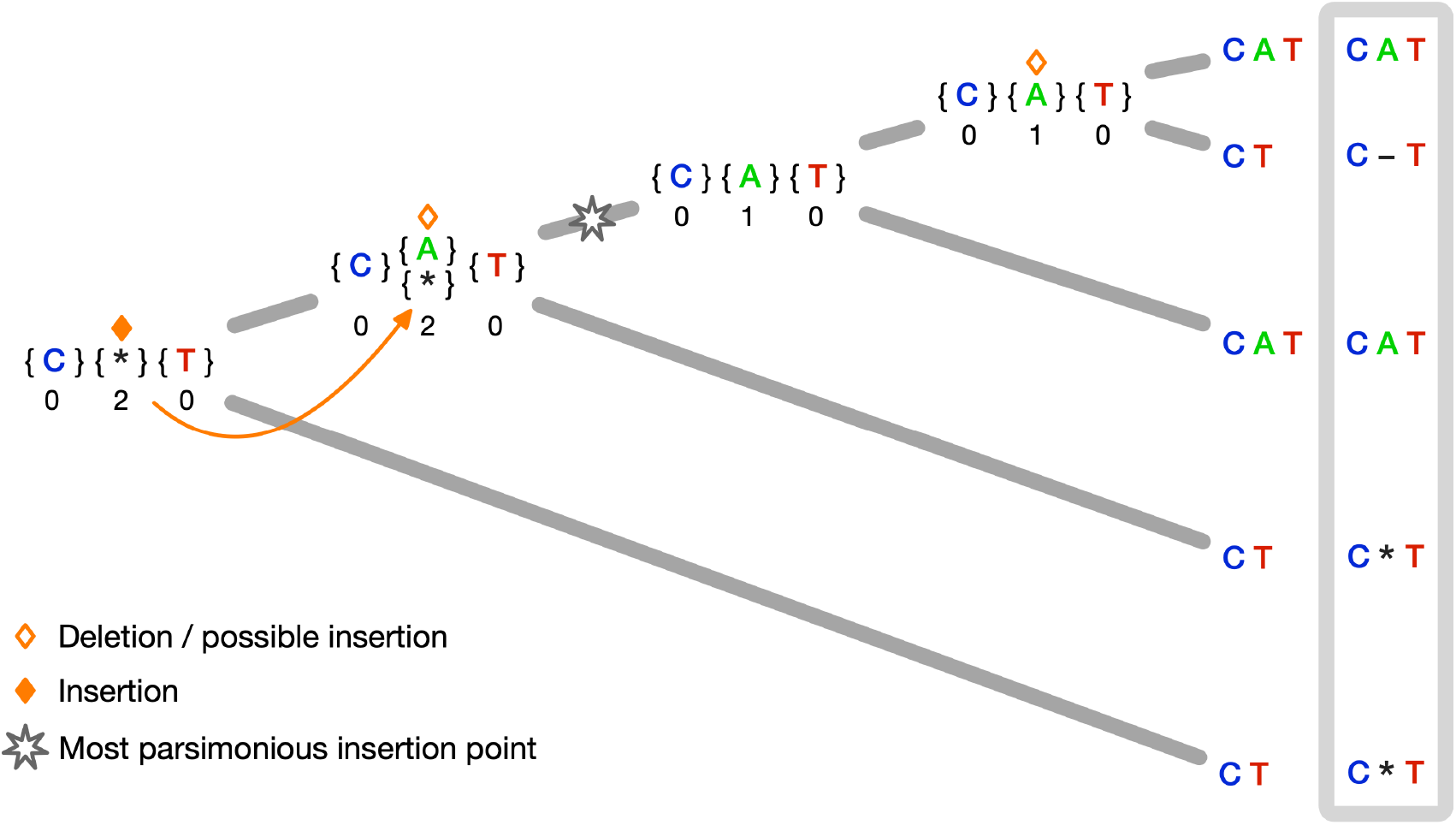
Multiple sequence alignment under indel-aware parsimony for a fixed guide tree. The parsimony scores per site are depicted under the respective parsimony sets. The middle site is marked as a deletion in the first internal node in the post-order tree traversal. In the subsequent alignment step, the column is aligned with a column containing a residue and the mark is removed. One step closer to the root, the site is again marked as a deletion and then as an insertion at the root node. Hence we change the internal reconstruction for the child node. The Dollo parsimony insertion point can be identified, and we can distinguish between gap characters representing deletions and placeholders.

## 3 Results

### 3.1 Validation on Simulated Data with Indels

To validate our methods, we used protein sequence datasets simulated with INDELible (Fletcher and Yang, 2009) under 54 parameter combinations, intended to imitate datasets with varying difficulty for phylogenetic inference. We simulated 16, 32 and 64 taxon trees with increasing divergence (represented by tree heights) of 0.8, 1.2, 1.7 and sampling fractions of 0.01, 0.25 and 0.99. Sampling fractions affect the tree topology: lower values yield longer external than internal branches, while higher values yield short external branches and longer internal branches (see Fig. 3). Amino acid sequences were simulated along the trees according to the WAG model (Whelan and Goldman, 2001) with indel rates of 0.01 and 0.05. For an overview of the simulation parameters and a detailed description, see Section 5.5.

**Figure 3.**
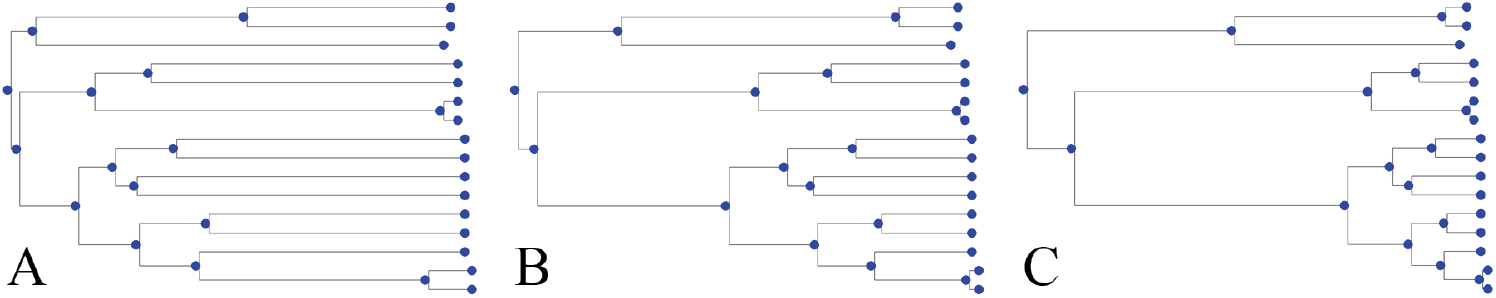
The figure shows the effect of different sampling fractions on tree topology for a 16-taxa tree. A low sampling fraction of 0.01 yields longer external branches (A), while higher sampling fractions of 0.25 (B) and of 0.99 (C) yield longer internal and shorter external branches.

#### 3.1.1 Performance of indelMaP for ancestral reconstruction of simulated protein data with long indels

Given the true tree and MSA, we assessed the reconstruction accuracy (see Section 5.7) of indelMaP against two parsimony methods: a conventional parsimony method, which treats gaps as an additional character, and GRASP, as well as against likelihood-based methods FastML, ARPIP and Historian (see Section 5.6). The overall character reconstruction accuracy for tree height 0.8 is summarised in Fig. 4. The overall accuracy of indelMaP was higher than that of the conventional parsimony for all parameter combinations. This improvement was larger for the higher indel rate, where indelMaP scored similar to or better than ARPIP. On the lower indel rate indelMaP is almost always outscored by Historian, while indelMaP scores better than Historian for the higher indel rate. Nevertheless, GRASP and FastML scored the best for all parameter combinations.

**Figure 4.**
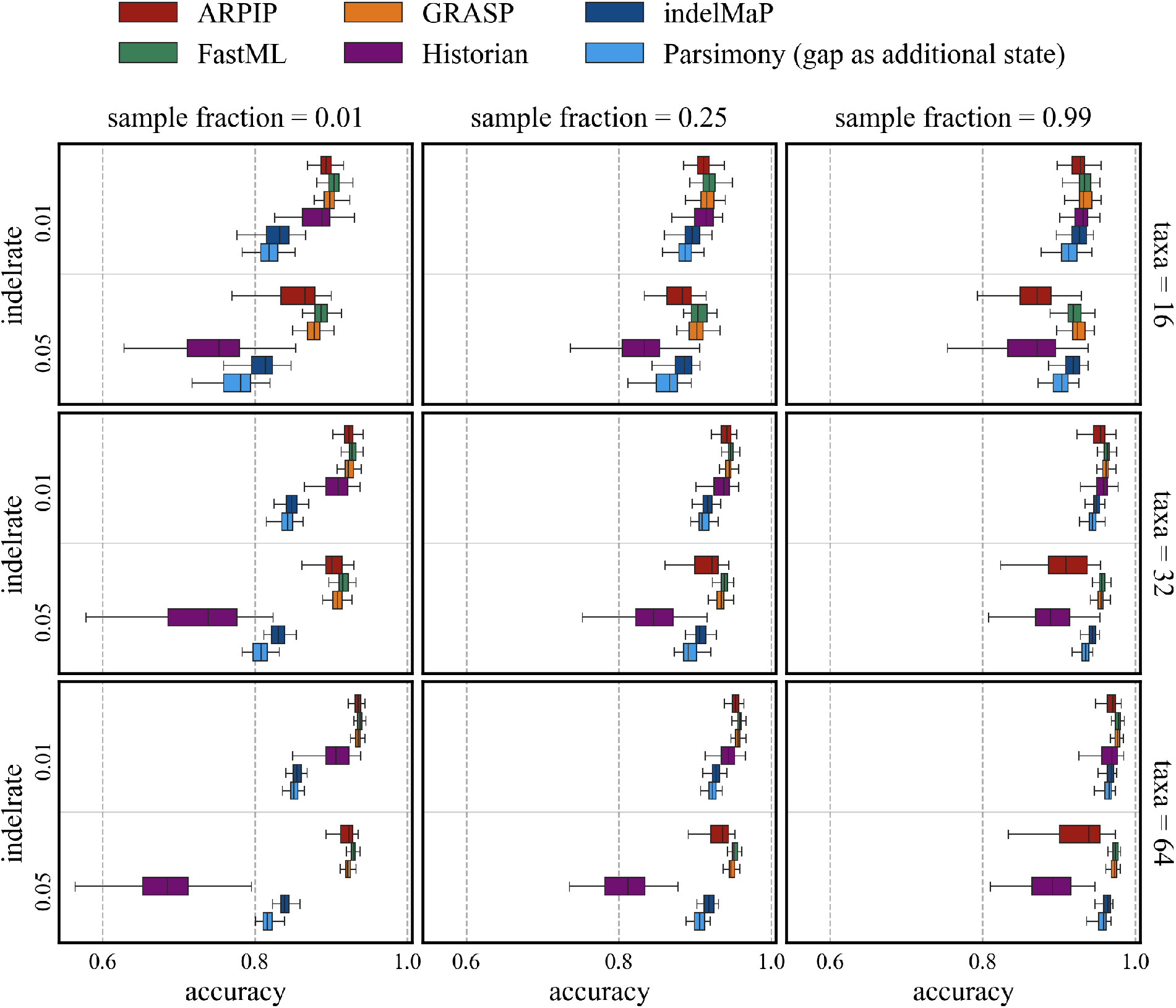
Overall character reconstruction accuracy for tree height 0.8.

In Fig. 5, the errors contributing to the overall accuracy are presented for an example of 16 taxa trees with a tree height of 0.8. For detailed results for all numbers of taxa, refer to Appendix A.2. The substitution error was the most significant for all methods, with indelMaP and conventional parsimony method having the highest. For the deletion error, indelMaP reaches quite a low error but not as low as FastML and was sometimes outscored by GRASP for the lower indel rate. ARPIP and indelMaP reach the lowest insertion error with zero or almost zero for all parameter combinations. Historian has the highest insertion error out of all methods.

**Figure 5.**
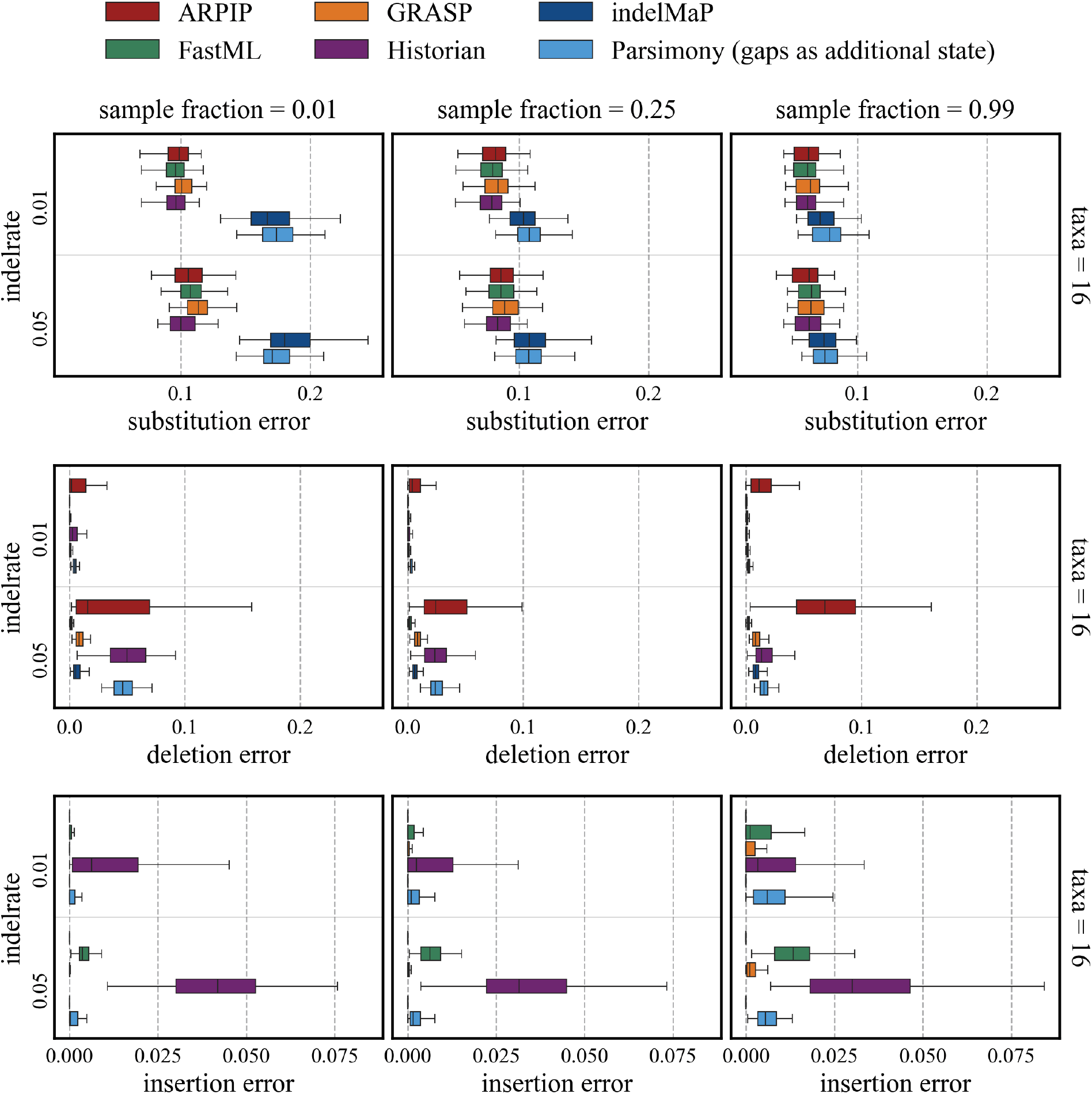
Substitution, insertion and deletion error for 16 taxa trees with tree height 0.8

For trees with a tree height of 1.2 and 1.7 the ordering of performances remains constant between the different methods; however, the overall performance declines (see Appendix A.1, Appendix A.3 and Appendix A.4).

indelMaP reaches the best reconstruction accuracy on sets with the highest sample fraction. To investigate if reconstruction accuracy and the node’s distance to the child nodes are related, we correlated substitution error and deletion error with the added distance to the child nodes of each node (see A.5 Figure Figs. SM 12 to SM 15. The insertion error is almost zero for all data sets and could not be correlated. We found a moderate correlation for the deletion error on sets with 0.99 sample fraction and 16 taxa and a strong positive correlation between substitution error and added distance to the child node. Additionally, the maximum substitution and deletion errors for one node were found for data sets with the highest sample fraction.

#### 3.1.2 Performance of indelMaP for multiple sequence alignment of simulated protein data with long indels

To validate the alignment quality of indelMaP on the simulated data, we evaluated estimated MSAs according to the Sum-of-Pairs Score (SPS) and Total Column Score (TCS) and according to its ability to infer indel rates and events during alignment estimation. For a detailed description of the quality measures, see Section 5.7) Additionally, we compared SPSs and TCSs obtained from indelMaP with those obtained from other intel-aware alignment methods: ProPIP, PRANK +F, Historian and the widely used aligner MAFFT, which is not indel-aware. First all MSA methods were given the true tree as the guide tree for MSA estimation. All MSA methods performed better with an increasing number of taxa and the higher sample fraction, with PRANK +F outscoring all other methods on all parameter combinations. The results for tree height 0.8 are summarised in Figure 6. Here, indelMaP reached SPS values close to PRANK +F and Historian on data sets with a lower indel rate, and outperformed MAFFT on the same data except for a sample fraction of 0.01. On the data sets with indel rate 0.05 indelMaP can only reach high scores on the set with sample fraction 0.99. Regarding TCS, indelMaP outscored MAFFT on all parameter combinations, except again for parameter combinations with the lowest sampling fraction. While SPS of ARPIP and indelMaP were comparable, indelMaP reached higher TCS values on almost all parameter combinations.

**Figure 6.**
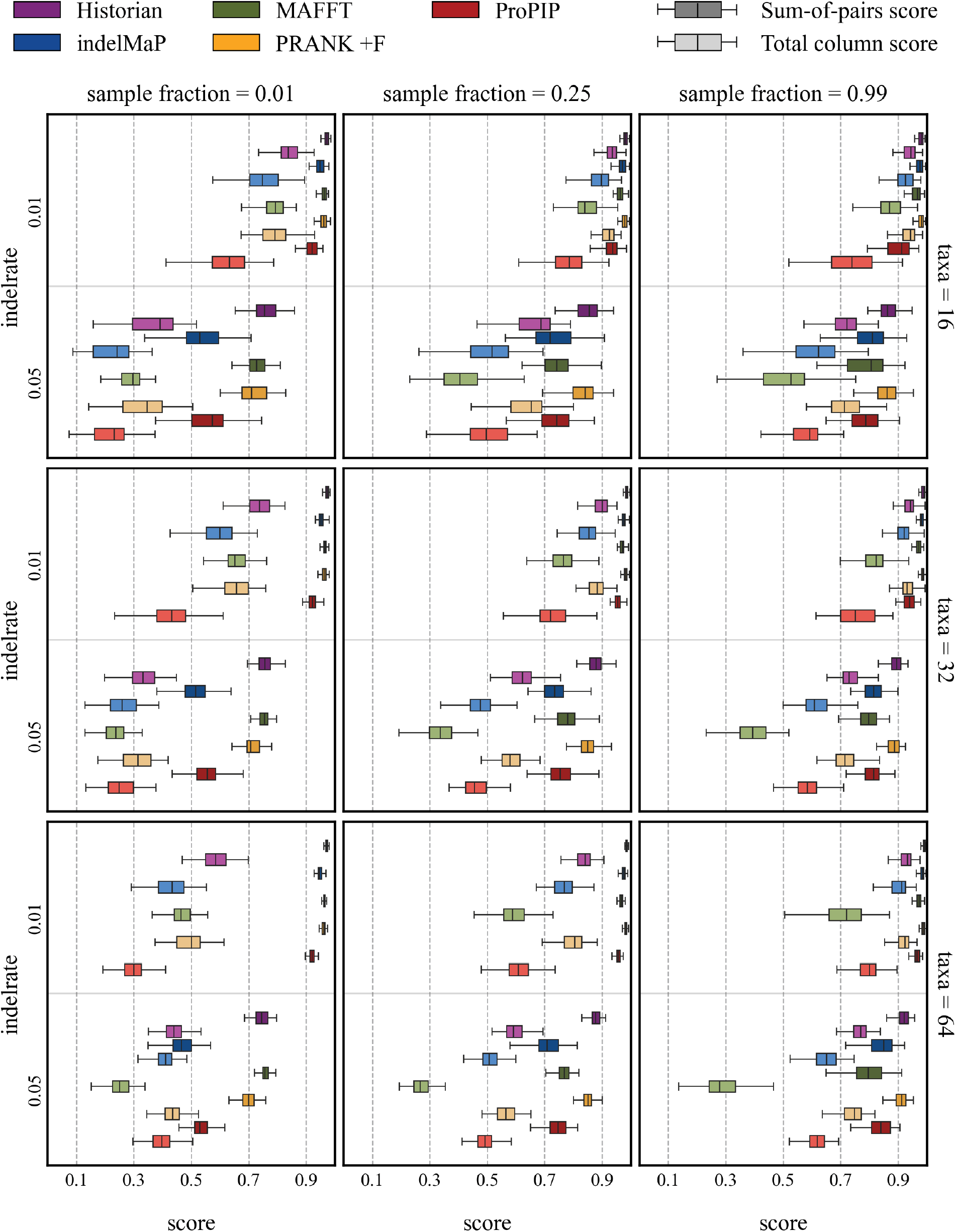
SPSs and TCSs scores for all parameter combinations with tree height 0.8.

We have observed that for trees with a height of 1.2 and 1.7, the ordering of SPSs and TCSs scores remains consistent across different methods. However, the overall performance declines. Please refer to Appendix A.6 for more information.

We evaluated indelMaP’s ability to estimate separate insertion and deletion rates and determine how well it recovers indel events during MSA estimation. We compared the relative rate (i.e., the estimated rate divided by the true rate) (Figure 7) and indel events (Figure 8) between indelMaP, Historian, and PRANK +F. All three are indel-aware, and Historian and PRANK +F have the highest MSA estimation scores. For indelMaP and Historian, we used the ancestral reconstruction given by the method. For PRANK +F, we used the events and ancestors given by PRANK +F and additionally reconstructed ancestors with indelMaP. IndelMaP tends to overestimate insertion rates compared to Historian. In contrast, deletion rates are close to the simulated rate except for the datasets with the lowest sample fraction. If indel rates are inferred from the reconstruction given by PRANK +F, insertion and deletion rates are assumed too high; even on the set with the highest sample fraction, we get indel rates that are ten to twenty magnitudes higher than the simulated rate. If we reconstruct ancestors with indelMaP based on the PRANK +F alignment, the rate estimates become close to the simulated rates on data sets with moderate to high sample fractions. Similar to the rates based only on indelMaP, insertion rates tend to be overestimated. For results for tree height 1.2 and 1.7 see A.6 Figure SM 18 and SM 19.

**Figure 7.**
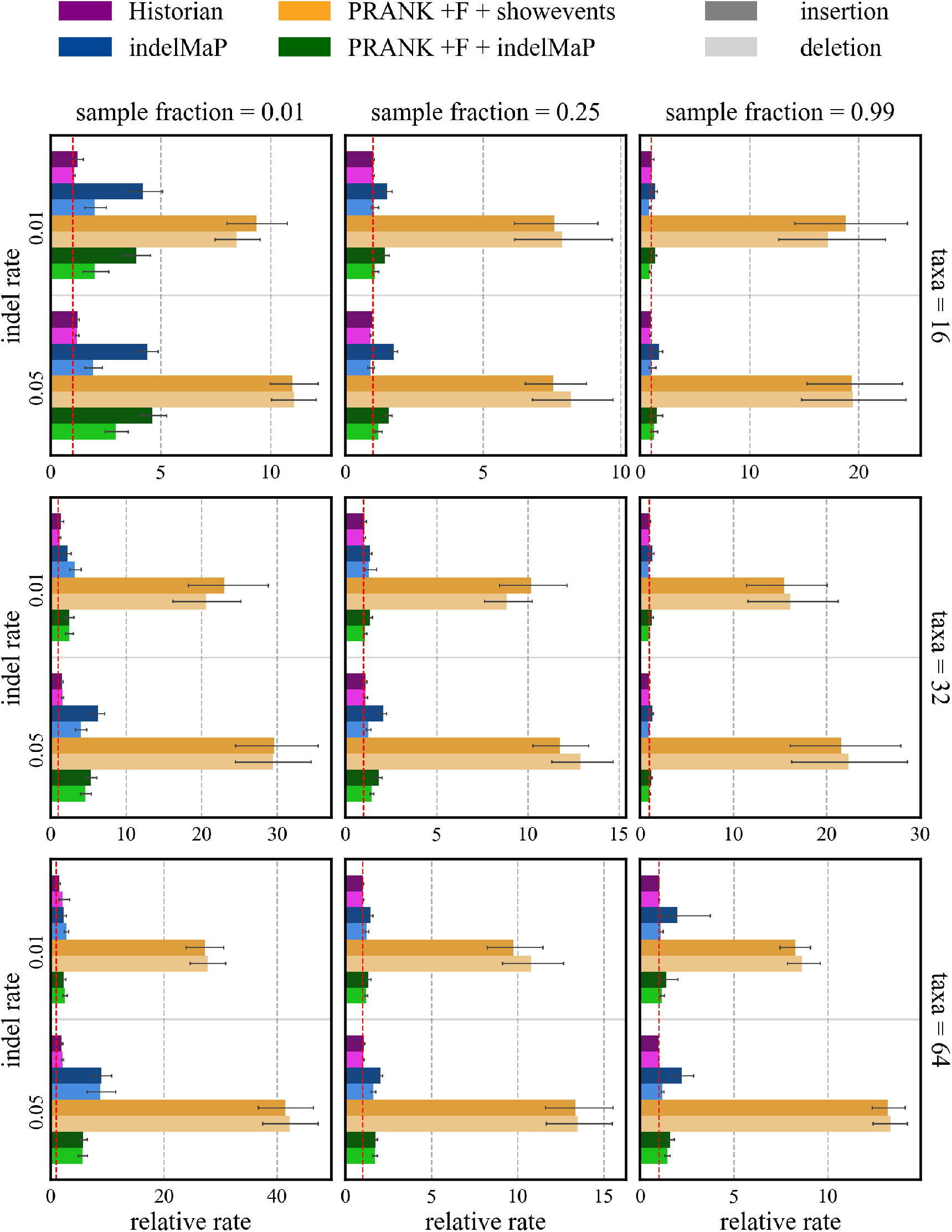
Relative indel rates, i.e. estimated rate over simulated rate, inferred based on Historian, indelMaP and PRANK +F. For PRANK +F we used the events given by the tool and additionally reconstructed ancestors with indelMaP based on the PRANK +F alignment. All tools were given the true tree as guide tree.

**Figure 8.**
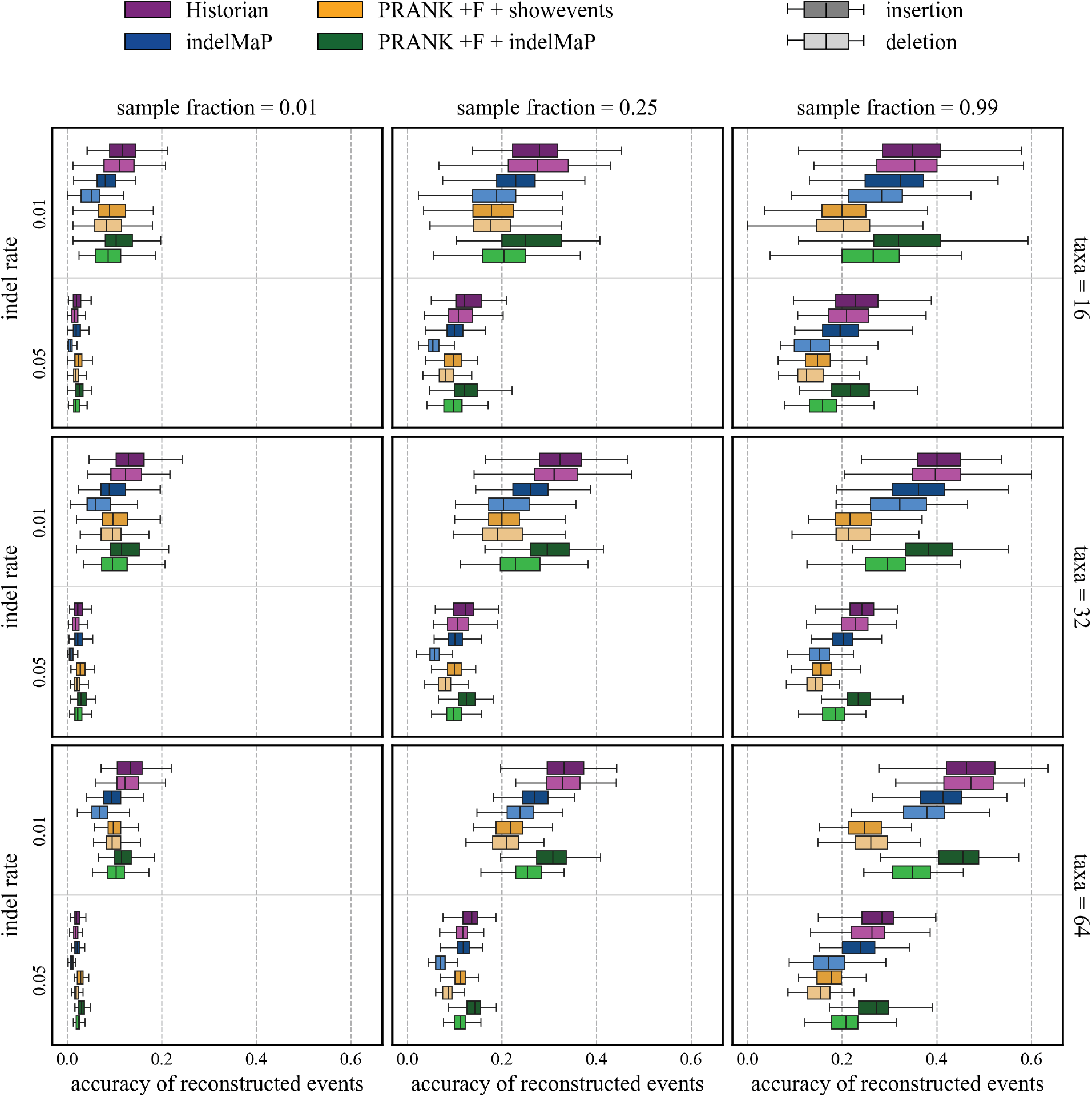
Proportion of accurate inferred insertion and deletion events for all parameter combinations with tree height 0.8. For PRANK +F we used the events given by the tool and additionally reconstructed ancestors with indelMaP based on the PRANK +F alignment.

**Figure 9.**
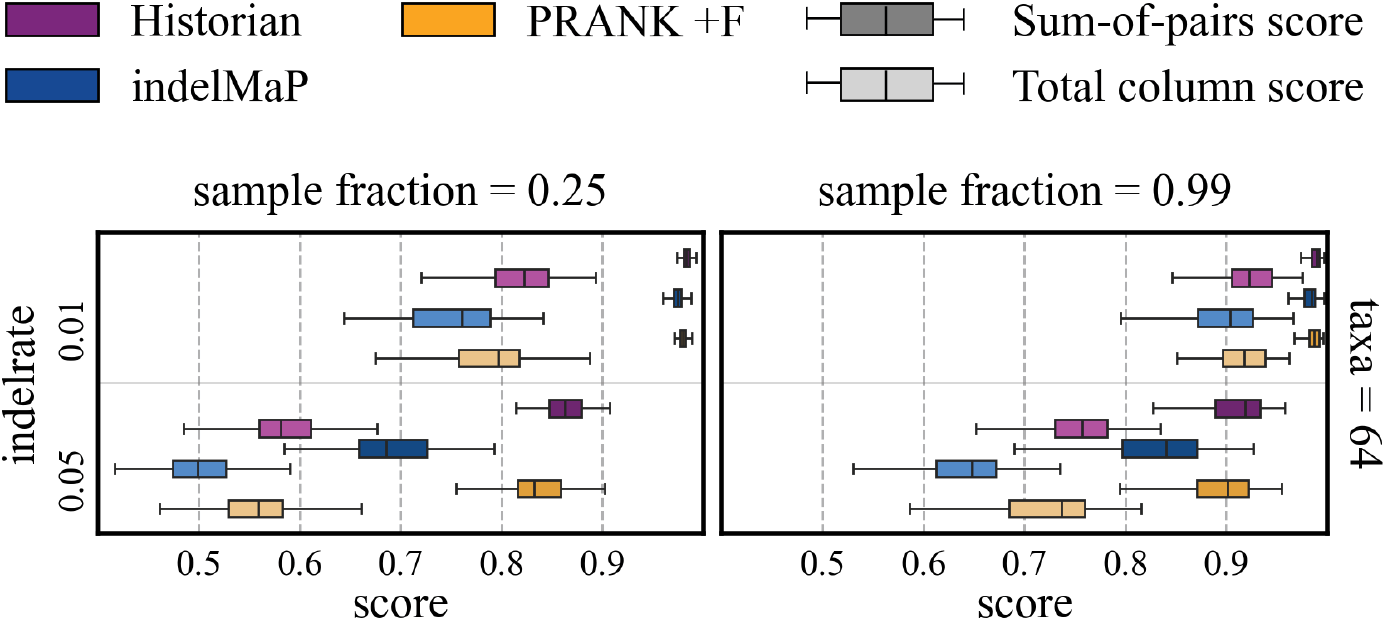
SPSs and TCSs scores for a subset of parameter combinations with tree height 0.8 and 64 taxa. All methods received the same estimated guide tree as guide tree for alignment.

**Figure 10.**
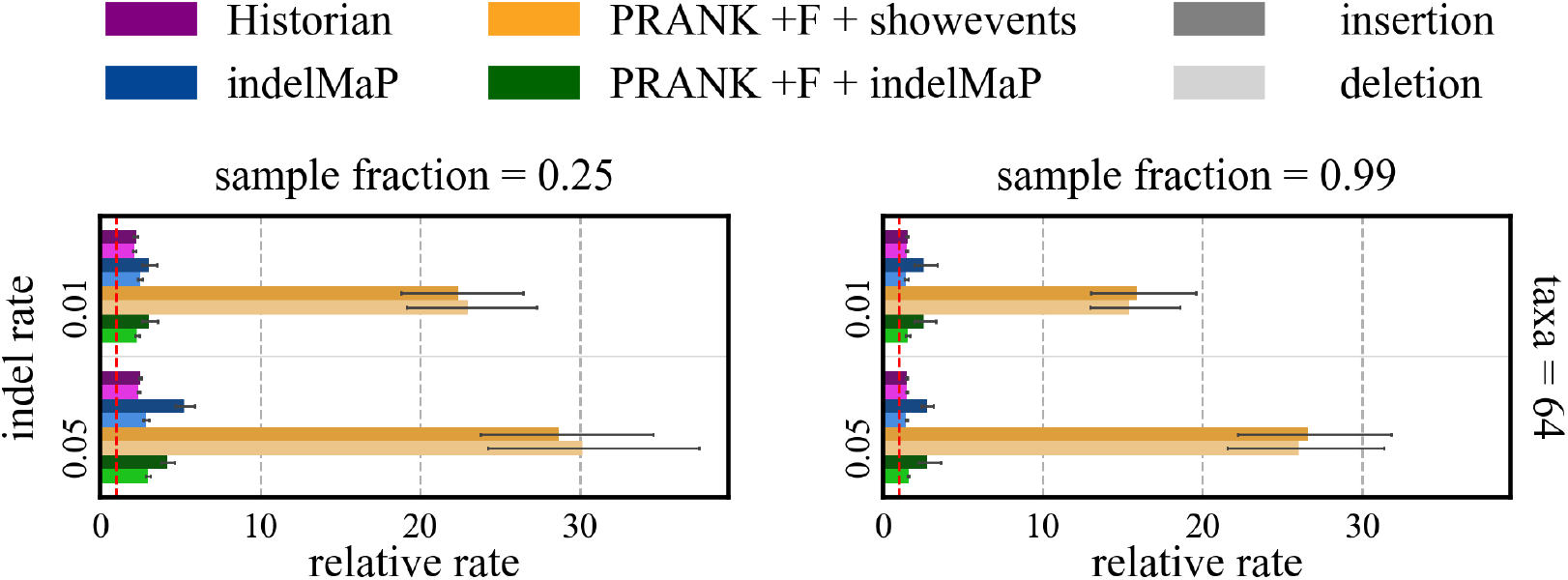
Relative indel rates, i.e. estimated rate over simulated rate, inferred based on Historian, indelMaP and PRANK +F. For PRANK +F we used the events given by the tool and additionally reconstructed ancestors with indelMaP based on the PRANK +F alignment. All tools were given an estimated tree.

**Figure 11.**
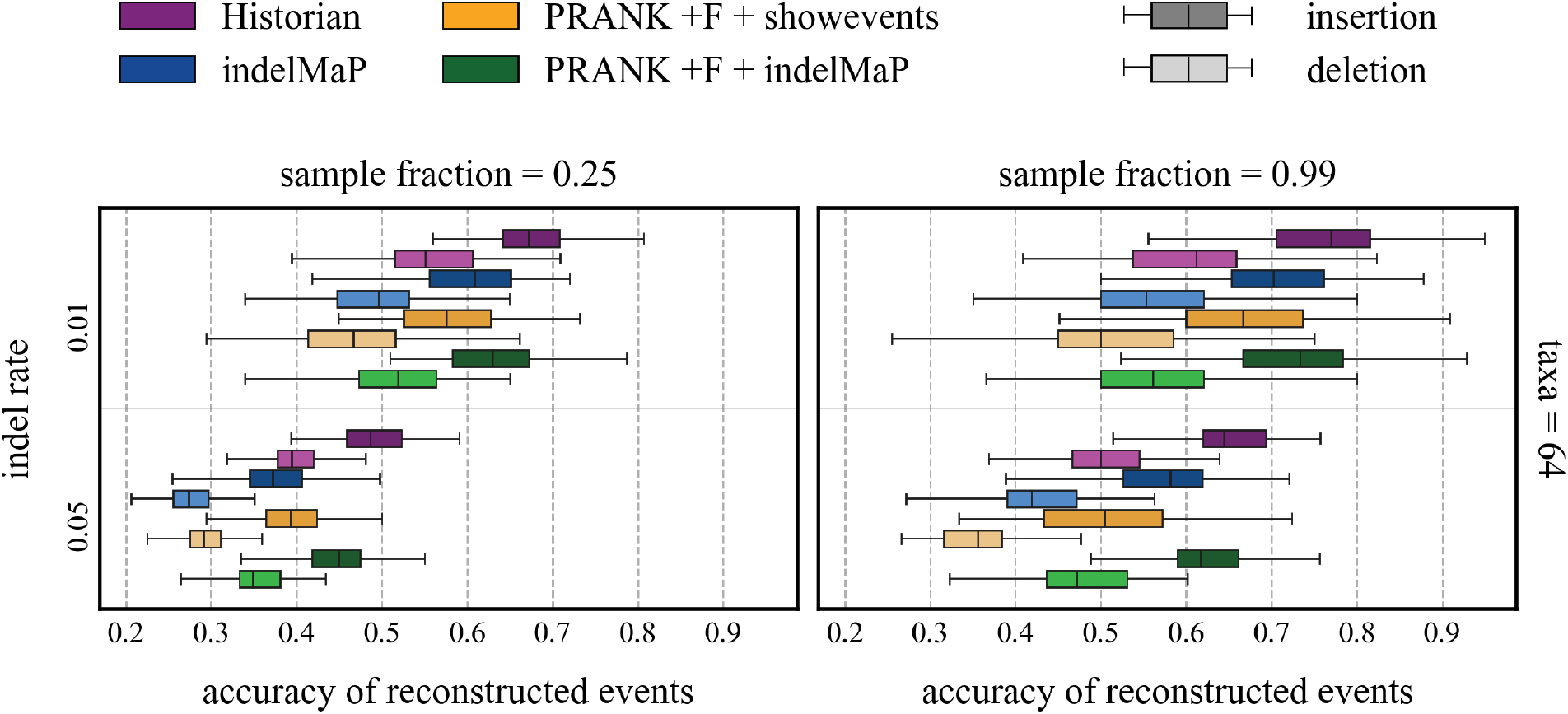
Proportion of accurate inferred insertion and deletion events. For PRANK +F we used the events given by the tool and additionally reconstructed ancestors with indelMaP based on the PRANK +F alignment. All tools were given an estimated guide tree.

Historian reaches the highest proportion of correctly estimated events on all data sets. IndelMaP reaches comparable accuracy for the low indel rate on the highest sample fraction. The quality declines for sets with the higher indel rate, especially for estimated deletion events. PRANK +F, if evaluated with the ancestral reconstruction from the tool itself, performs worst on the highest sample fraction. For the indelMaP reconstruction based on a PRANK + F alignment, PRANK +F reaches better accuracy, sometimes outscoring indelMaP for accurately inferred insertion events and on the higher indel rate for accurately inferred deletion rate. For results for tree height 1.2 and 1.7 see A.6 Figure SM 20 and SM 21.

Next, we chose PRANK +F, Historian and indelMaP for further validation to explore the effect of an estimated guide tree on MSA estimation. We let PRANK +F run without specifying a guide tree and then used the estimated tree as input for indelMaP and Historian. The evaluation of SPS and TCS for a subset of parameter combinations is shown in 9. While indelMaP can reach comparable SPS and TCS scores to PRANK +F and Historian on the lower indel rate, it falls behind for the higher indel rate, especially in the less densely sampled data set.

Furthermore, we investigated the effect of an estimated guide tree on the inferred indel rate 10 and proportion of accurately reconstructed events 11 on the smaller subset. While all tools overestimate insertion and deletion rates on the less densely sampled data set, indelMaP shows a bias towards insertions. Historian also has the highest proportion of accurately inferred events, closely followed by indelMaP and PRANK +F with an indelMaP reconstruction. For the closely sampled set, Historian infers rates that almost perfectly match the simulated rate, while indelMaP slightly overestimates insertion rates and infers deletion rates, which are very close to the true rate. This also holds for estimates based on PRANK +F alignment and an indelMaP reconstruction. The ordering for the proportion of the correctly inferred events is the same as for the less closely sampled data set but with higher proportions. Notably, PRANK +F with indel rates based on their reconstruction produces indel rates several magnitudes higher than the simulated rate.

#### 3.1.3 Sensitivity analysis for gap opening and gap extension factor

We refrained from tuning the gap opening and gap extension penalty for the affine gap score to the specific benchmark and chose a gap opening penalty factor of 2.5 and a gap extension factor of 0.5 based on recommendations by Altschul et al. (Altschul and Erickson, 1986), both are multiplied by the average substitution rate. To investigate the sensitivity of the method to gap opening and gap extension factor we conducted a small sensitivity analysis on a subset of the data. The parameters we chose are sets with tree height 0.8, sample fraction 0.25 and 0.99, indel rate 0.01 and 0.05 and 64 taxa. We evaluated the quality of the estimated alignment based on SPS and TCS. The results for SPS are shown in Figure 12 and for TCS in Figure 13. For the lower indel rate SPS and TCS are not very sensitive, SPS and TCS reach their maximum for gap opening factors larger than one and for gap extension factors larger than 0.3. For the higher indel rate the alignment quality is more sensitive to gap opening and gap extension factor. The SPS and TCS reach their maximum for an gap opening factor of 1.5 and gap extension factors of 0.5 and 0.7.

**Figure 12.**
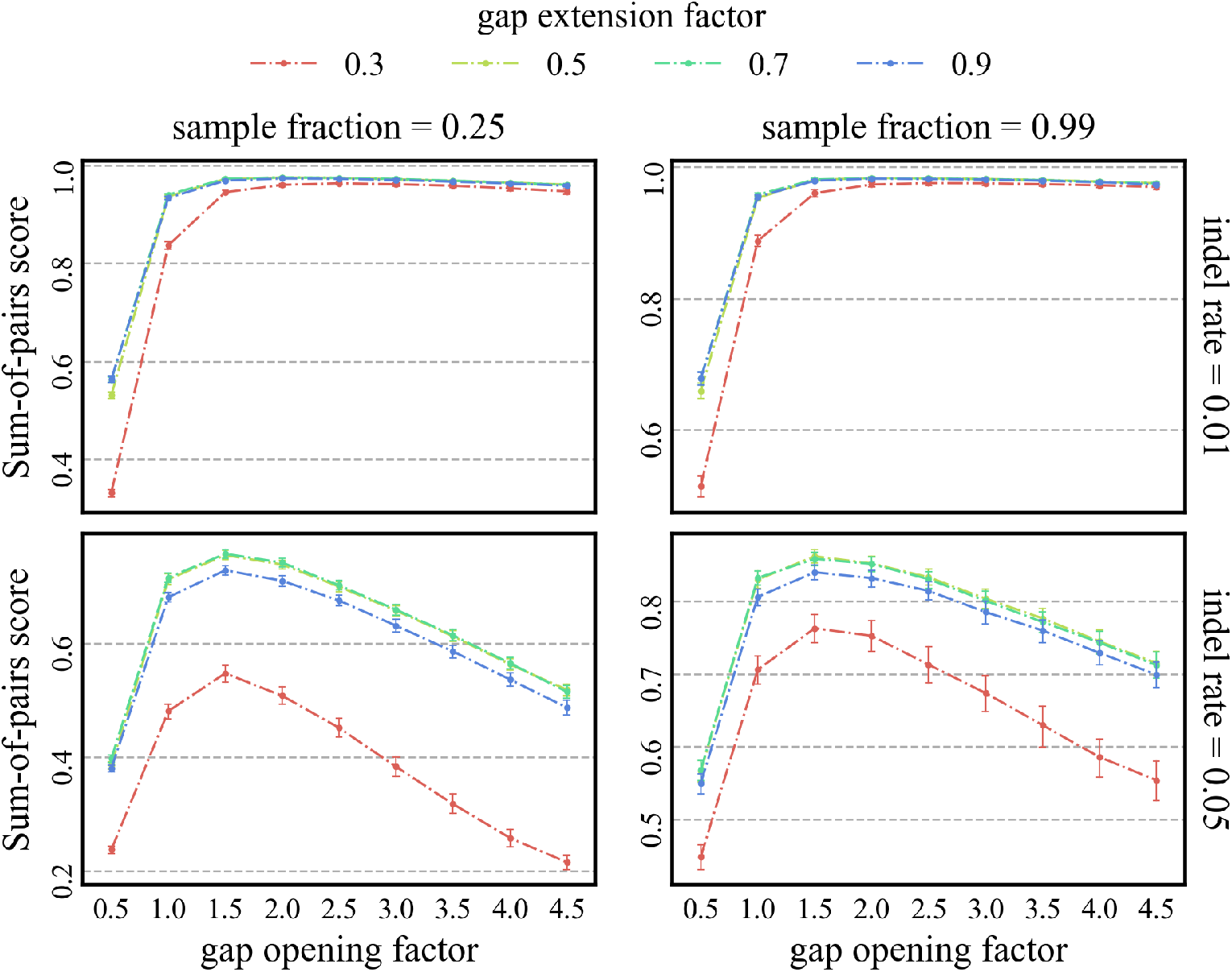
Effect of different combinations of gap opening factor and gap extension factor on the Sum-of-pairs score.

**Figure 13.**
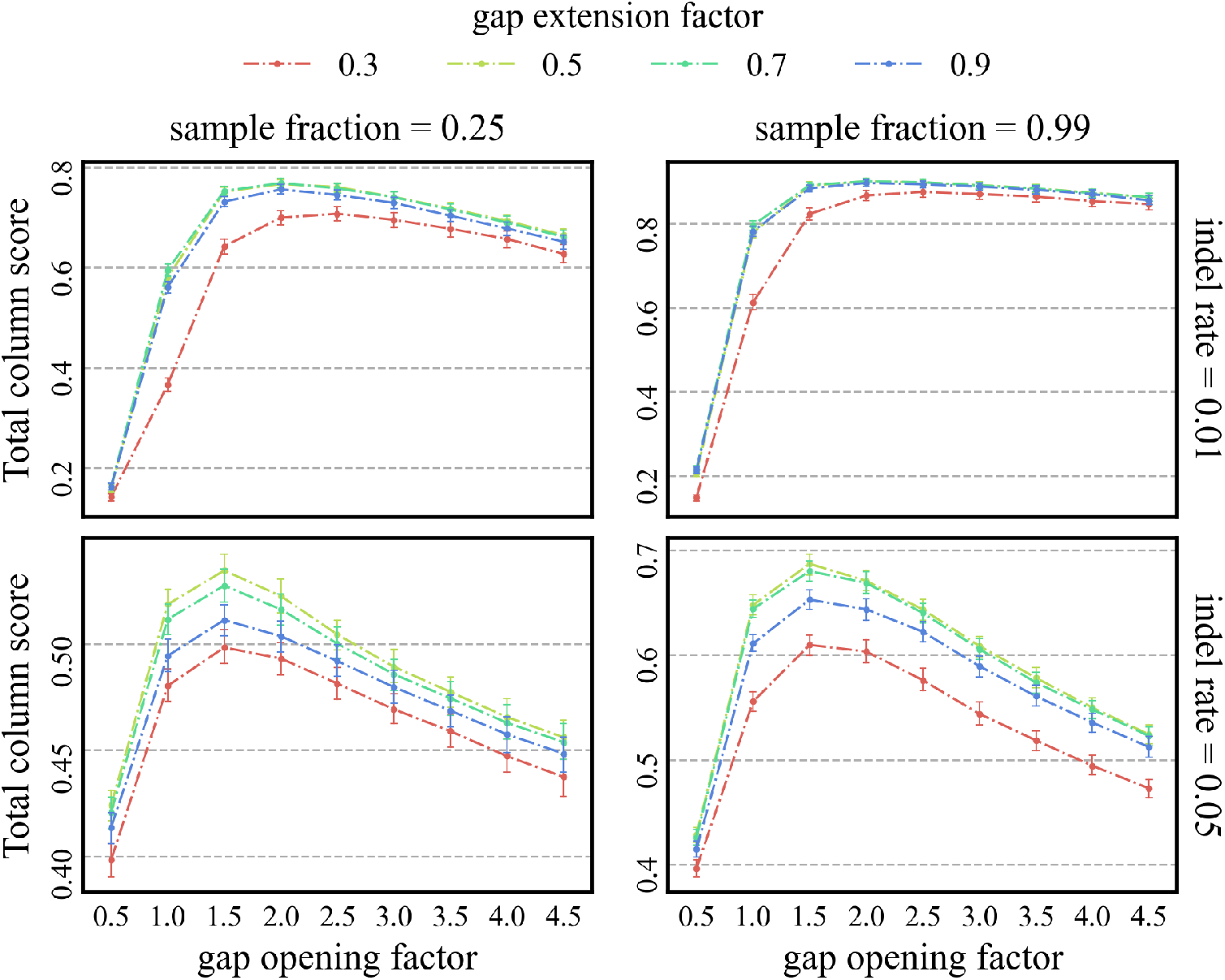
Effect of different combinations of gap opening factor and gap extension factor on the Total column score.

#### 3.1.4 Time benchmark for multiple sequence alignment and ancestral reconstruction with indelMaP

To demonstrate the efficiency of our method across diverse parameter combinations, we expanded our simulation study to include large datasets with 200 taxa. We also included one set with 400 and 800 sequences and benchmarked the faster methods on that set. Results can be seen in A.7 Figure SM 22 for MSA and A.7 Figure SM 24 for ASR estimation. We benchmarked all MSA and ASR methods using the hyperfine tool on a quiet machine with a 2.3 GHz Quad-Core Intel Core i7 processor and 16 GB RAM. The results for MSA estimations for tree height 0.8 are shown in Figure 14. The results revealed that the Python implementation of indelMaP exhibited elevated computational times, rendering it impractical to analyse exceptionally large datasets. While its processing speed surpasses that of MSA estimation under PIP, there were instances where it trailed behind PRANK and displayed a notable difference compared to MAFFT and Historian. To address this issue, we re-implemented the method in Rust, which significantly improved computational efficiency. This optimised version of indelMaP not only outperformed its Python counterpart but also demonstrated a remarkable speed, significantly surpassing MAFFT for parameter combinations with a high indel rate. Historian’s computational time remains low for datasets up to 200 sequences but becomes relatively slow for larger sets.

**Figure 14.**
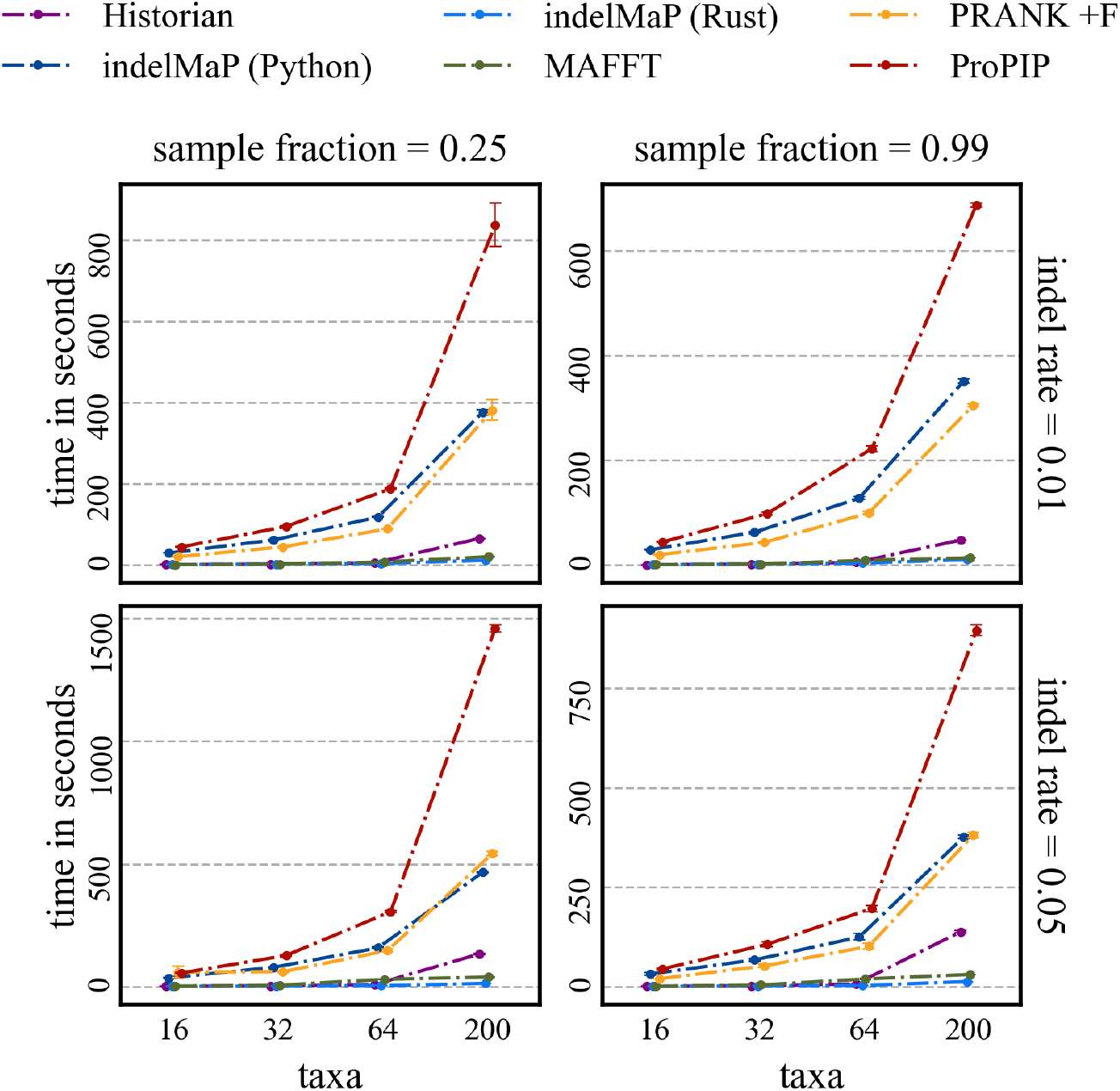
Computational time benchmark of MSA methods, under various parameter combinations with tree height 0.8.

Compared to MSA estimation, ASR inference inherently imposes a relatively lower computational burden. While the Python implementation of indelMaP has superior speed to methods like ARPIP and FastML, it exhibits a similar or slightly slower computational time than GRASP, Historian and the conventional parsimony method. However, indelMaP already demonstrates sufficient efficiency for handling large datasets. Nevertheless, the prospect of a Rust implementation for indelMaP remains promising, with the potential to further enhance its processing speed. The results are summarised in A.7 Figure SM 23 for tree height 0.8. For data sets with 400 and 800 taxa trees, the computation times of Historian and indelMaP remain relatively fast. GRASP, which is also still fast for 400 taxa trees, becomes intractable for 800 taxa trees.

#### 3.1.5 IndelMaP as a computational approximation for the likelihood score under PIP

Methods using an explicit indel model can be computationally expensive. To investigate if indelMaP could be used as a computational approximation for phylogenetic methods using the PIP model, we calculated the indelMaP score and conventional parsimony score treating the gap character as an additional state for the true tree and MSAs estimated by ProPIP. Since PIP reconstructs single residue indel events, we used a linear gap cost for indelMaP. The Pearson correlations for all parameter combinations with tree height 0.8 are summarised in Table 1. For the results for tree height 1.2, see Appendix A Table SM 1. IndelMaP reaches a higher negative correlation with the log-likelihood score on all parameter combinations than the conventional parsimony score. On 64 taxa trees, the indelMaP score is almost perfectly negatively correlated with the log-likelihood score under PIP.

**Table 1:** Pearson correlation between the log-likelihood score under PIP (log LK), indelMaP and the conventional parsimony score treating the gap character as an additional state (MP). The upper triangular matrix shows correlations for the indel rate of 0.01 and the lower for the indel rate of 0.05. All parameter combinations are for tree height 0.8.

|  | 16 taxa |  |  | 32 taxa |  |  | 64 taxa |  |  |  |
| --- | --- | --- | --- | --- | --- | --- | --- | --- | --- | --- |
| log LK | - | -0.989 | -0.979 | - | -0.983 | -0.979 | - | -0.995 | -0.992 | 0.25 |
| indelMaP | -0.997 | - | 0.995 | -0.994 | - | 0.98 | -0.996 | - | 0.994 |  |
| MP | -0.975 | 0.995 | - | -0.982 | 0.99 | - | -0.99 | 0.995 | - |  |
| log LK | - | -0.933 | -0.894 | - | -0.945 | -0.916 | - | -0.997 | -0.995 | 0.99 |
| indelMaP | -0.973 | - | 0.982 | -0.979 | - | 0.992 | -0.997 | - | 0.995 |  |
| MP | -0.952 | 0.983 | - | -0.958 | 0.991 | - | -0.992 | 0.996 | - |  |
|  | log LK | indelMaP | MP | log LK | indelMaP | MP | log LK | indelMaP | MP |  |

### 3.2 Indel Pattern Analysis of the HIV-1 Envelope Protein gp120

We tested our approach by analysing the between host indel diversity in the variable regions of the envelope glycoprotein gp120. The data set and methods are detailed in Section 5.8. The variable regions of the envelope glycoprotein protein (gp) 120 are prone to indel events, which are driven by natural selection and closely linked to the fitness of the virus (Wood et al., 2009). While the length variability of gp120 is known to contribute to the virus’s immune escape, the specific indel patterns and rates responsible for this mechanism have only recently gained attention (Palmer and Poon, 2019). As gp120 is responsible for binding to the host cell, it is a primary target for vaccine development. Therefore, understanding indel patterns in gp120 and reconstructing ancestral proteins, including indels, will be crucial for advancing successful immunisation strategies (Pantophlet and Burton, 2006). Figure 15 (top) shows the mean fraction of inserted or deleted sites per variable region per year compared to the whole gp120 sequence. The variable region V3 accumulated the least insertion and deletions. Region V5 accumulated the most inserted sites and V4 the most deletions. In Figure 15 (bottom), the indel length frequencies of the different regions are depicted. While the median deletion length in V3 was one residue, the same as in the whole sequence of gp120, the median deletion length was two for the other variable regions. V1 had a median of four and V3 of three for insertion lengths compared to two for the overall sequence.

**Figure 15.**
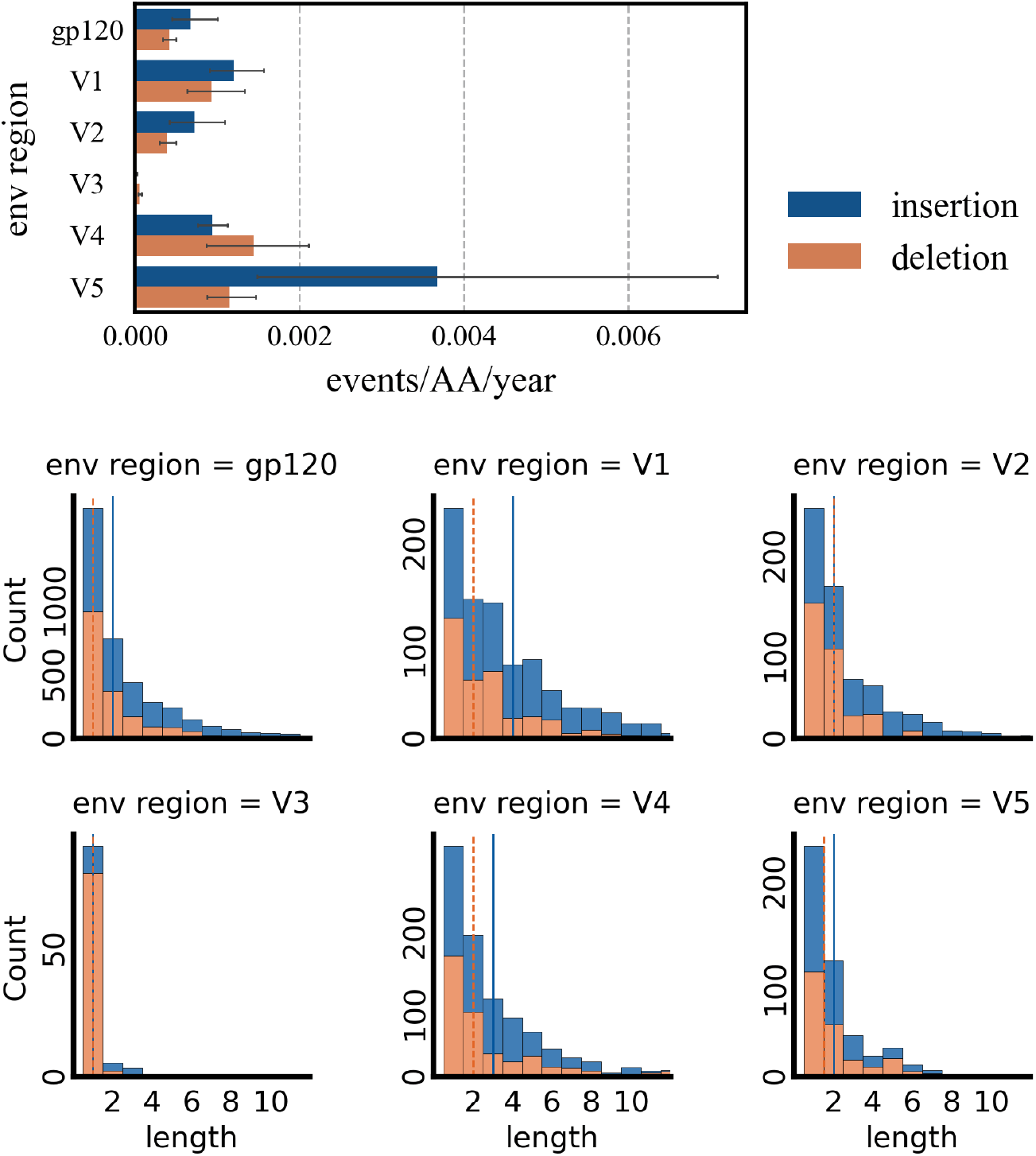
The top figure shows the mean fraction of insertion or deletion events over the length of the region per year for the whole gp120 sequence and the variable regions V1-V5. Fliers indicate the 95th confidence interval of the estimate. On the bottom, the insertion and deletion frequencies per region are shown. The vertical lines indicate the median length for insertion or deletion lengths. Indels with a length over 11 residues are not pictured.

## 4 Discussion

Our results underscore the effectiveness of the indelMaP method in addressing the challenges posed by insertion and deletion events in evolutionary analyses. For ASR, indelMaP reaches comparable overall reconstruction accuracy to other state-of-the-art indel-aware methods and consistently outscores the standard parsimony that treats a gap character as an additional state. Notably, indelMaP displays the lowest cumulative indel error among all considered approaches, even on sets with distantly related taxa, highlighting the robustness of our new method in dealing with indel events and its potential for more accurate indel reconstructions in analyses where indels play a significant role.

We observed that more sophisticated methods tend to achieve lower substitution error rates overall, with indelMaP’s performance showing a decline on distant datasets with fewer taxa. Furthermore, the strong correlation between substitution error and a node’s distance to its child nodes, even in datasets with a high sample fraction, suggests that indelMaP’s reconstruction accuracy for substitution events diminishes for deeper nodes. This observation aligns with the complexity inherent in distant relationships, where the incorporation of branch-length information in probabilistic methods enhances reconstruction accuracy.

Similarly, indelMaP alignment quality is comparable to probabilistic methods on densely sampled datasets, although it falls behind on sparse and distant datasets. Additionally, it is important to note that indelMaP exhibits a bias towards insertions during alignment estimation, particularly if the guide tree is estimated. While indelMaP may achieve a lower accuracy for indel rate estimation and event reconstruction than Historian, it does surpass PRANK +F in this regard. Furthermore, employing indelMaP as a reconstruction method on alignments generated by PRANK +F proves advantageous for indel pattern analysis. Although indelMaP alignments should be regarded as approximations due to their inherent biases, our sensitivity analysis for gap opening and extension factors showed that indelMaP’s alignment quality could be improved if gap penalties were optimised for the specific data set.

Overall, the results emphasise the suitability of parsimony-based approaches in scenarios where the relationships between taxa are relatively easy to establish, and the emphasis shifts towards minimising the associated computational burden. Our results show that indelMaP can align large datasets with remarkable speed, making it the perfect choice for epidemiological datasets to avoid downsampling and leverage the additional information of dense taxonomic sampling (DeWitt et al., 2018; Feng et al., 2020; de Abreu-Jr et al., 2020). Furthermore, the fast computational time, together with the almost perfect negative correlation between the indelMaP score and the log-likelihood score under PIP, demonstrates that indelMaP could be used as a computational approximation during the optimisation process in methods that use PIP as an explicit indel model.

Analysing indel patterns within the envelope protein gp120 highlights indelMaP’s capacity to offer novel insights into the evolutionary implications of insertion and deletion events within biologically significant sequences. In the past, these events have been studied over length variations (Zolla-Pazner and Cardozo, 2010) or as aggregated indel events (Palmer and Poon, 2019). However, leveraging an indel-aware approach allows for separating insertion and deletion events, shedding light on their distinct roles and contributions to the evolutionary dynamics. Our approach introduces a novel perspective on gp120, and our findings are consistent with previously reported length variability of the variable loops. While it was previously observed that regions V1 and V2 tend to grow in length in chronically infected patients (Sagar et al., 2006), it is essential to note that the reported higher insertion rate for V1, V2, and V5 could be overestimated due to the tool’s insertion bias. Moreover, the higher rate for V5 could also be inflated, as this region is very short, approximately 11 amino acids long, and challenging to extract accurately.

In summary, indelMaP presents a compelling alternative for treating gaps within parsimony-based approaches while accounting for the distinct evolutionary trajectories of insertion and deletion events. The method’s compatibility with affine gaps demonstrates its potential as a computational approximation not only for explicit models involving single residue indel events but also suggests its feasibility for long indel models, under which accurate MSA estimation remains challenging. Moreover, beyond its applicability for ASR and MSA, indelMaP could prove valuable for indel-aware tree search and within joint alignment and tree search algorithms.

## 5 Methods and Materials

**Notation**. The guide tree *τ* is given by a rooted binary phylogenetic tree, defined by a set 𝒱 of nodes, with *N* leaf nodes and a set ℰ *⊂ 𝒱×𝒱* of edges. Each leaf is associated with one of the sequences we want to align, given by a string of characters from a finite alphabet Σ, containing nucleotides or amino acids. For the alignment, we extend Σ with the gap character given by *ϵ*. Let *C ∈ ℕ*^|Σ|*×*|Σ|^ be an arbitrary symmetric cost matrix where *C*_*i,j*_ is the associated cost of substituting the character *i* with the character *j* and *a, b* be the penalty for opening and extending a gap, respectively. Consequently, a gap of length *k* has the cost *a* + *b*(*k −* 1).

### 5.1 Ancestral sequence reconstruction under indel-aware parsimony

The method is an extension of Fitch’s algorithm and reconstructs ancestral sequences for a fixed tree *τ* and alignment *A* and calculates the associated indel-aware parsimony score. Before we traverse each node *v* of *τ* in post-order to calculate the parsimony score, *s*_*k*_ and parsimony set 𝒫_*k*_ for each site *k* ∈ {1, …, |*A*|}, we determine Dollo parsimony insertion point of each extant residue. The method requires two Boolean vectors in the alignment length for each node. The vector *h* stores a true value if an insertion occurred between the particular node and one of its child nodes. The vector *g* stores the gaps in *A* if a deletion occurred above the current node or an insertion occurred below the current node. The algorithm must skip these sites when we want to determine if we are currently in an open gap.

#### Insertion points

We define each residue’s Dollo parsimony insertion point as the most recent common ancestor node of all leaves containing a residue at the site. All leaves outside the sub-tree defined by this insertion point will necessarily have a gap character at this site. Any leaves with a gap character within the sub-tree will have to be explained by deletion events. For column *k* in the alignment, we identify the leaves associated with a residue and traverse the tree post-order to find the most common ancestor *a ∈ 𝒱* of those leaves. Per definition, the insertion occurred along the branch from the parent node *pa*(*a*) to *a*. We flag the insertion point in node *pa*(*a*), i.e., *h*_*k*_ = 1. Note, if *a* is the root, no insertion happens inside the current tree *τ*, and all gap characters at the site are treated as deletions.

#### Post-order tree traversal

The algorithm traverses each node *v ∈ 𝒱* in post-order. If *v* is a leaf, we initialise the parsimony score *s*_*k*_ to zero, and the parsimony set 𝒫_*k*_ only holds the character present at site *k*. If *v* is an internal node with two child nodes *x* and *y*, we determine the parsimony score *s*_*k*_, the parsimony set 𝒫_*k*_ and the flag *g*_*k*_ for each site *k* ∈ *k* ∈ {1, …, |*A*|} over the parsimony sets 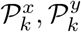 the insertion flag *h*_*k*_ and the vector *g* of the current node.

Suppose neither 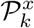 nor 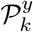 includes a gap character at the site. The site corresponds to a match or substitution, and the parsimony score is calculated according to the cost matrix. The new parsimony set is either given by the union or intersection of 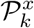 and 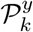 We set *g*_*k*_ = 0 since we do not skip the site to determine if we are currently in an open gap.

If 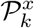 contains a gap character while 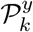 contains a residue, the site corresponds to an insertion or deletion. If *h*_*k*_ = 0, the site is a deletion, and 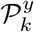 gives the new parsimony set. We use the entries of *g* and *h* before *k* to determine if we are currently in an open deletion. Let *l* = *k −* 1 while *g*_*l*_ = 1 or *h*_*l*_ = 1 set *l* = *l−* 1. If 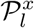 also contains a gap character and *g*_*l*_ = *h*_*l*_ = 0, we are extending a deletion in *k*. Otherwise, we are opening a new gap. We set *g*_*k*_ = 0 since we are not in an open insertion. If we flagged the site as an insertion with *h*_*k*_ = 1, we let the new parsimony set *𝒫*_*k*_ only include the gap character to ensure that the site is not penalised again in the ancestral nodes. To determine if we are currently in an open insertion, let *l* = *k −* 1 and set *l* = *l −* 1 while *h*_*l*_ = 0 and *g*_*l*_ = 1. If 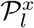 contains the gap character and *h*_*l*_ = 1, we extend an insertion in *k*. Otherwise, we are opening a new insertion. We set *g*_*k*_ = 1 since the site has to be skipped.

The symmetrical situation where 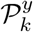 contains a gap character while 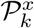 contains a residue is treated analogously.

If 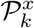 and 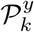 only include the gap character, either an insertion occurred in a descendant node or a deletion occurred in an ancestor node. Consequently, the event is penalised in a preceding or subsequent node, not the current node. The new parsimony set only includes the gap character and *g*_*k*_ = 1.

The computation of the score *s*_*k*_, *𝒫*_*k*_ and *g*_*k*_ is summarised in the following formula:

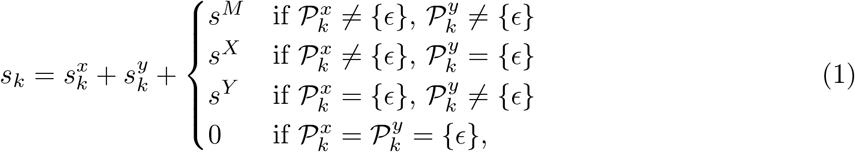

where

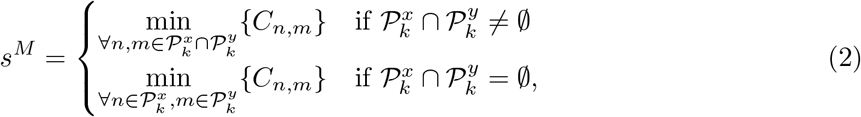

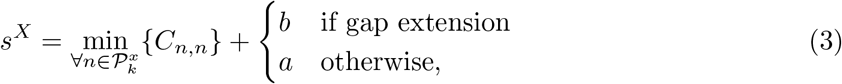

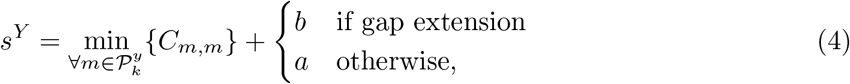

and

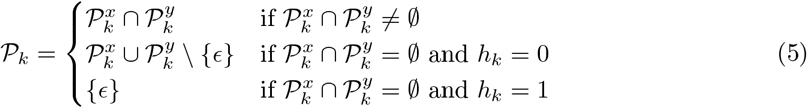

and

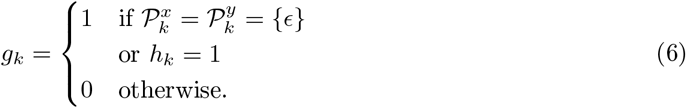

We determine the score of the whole tree *τ* by summing over all scores per site at the root node.

#### Pre-order tree traversal

The ancestral sequences at each internal node *v* ∈ 𝒱 are reconstructed during a pre-order tree traversal from the root to the leaves. Let 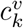 denote the most parsimonious reconstruction for node *v* at site *k*. We determine 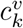 similar to Fitch’s algorithm. If *v* has a parent node *pa*(*v*) and 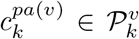 we set 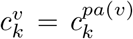. Otherwise, instead of making a random choice between characters, we assign the character in 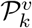 associated with the lowest cost to 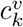.

### 5.2 Multiple sequence alignment under indel-aware parsimony

We propose a progressive alignment method that traverses a rooted binary guide tree in post-order. At every internal root, it aligns two sub-alignments according to a dynamic programming approach guided by the indel-aware parsimony criterion. During the post-order tree traversal, the most parsimonious site reconstructions for each internal state are memorised in parsimony sets. To identify the Dollo parsimony insertion point for each site and treat deletions and insertions as separate evolutionary events, we implemented an option to flag gap columns as proposed in (Löytynoja, 2014).

Here, we only describe the algorithm for affine gaps, which follows Gotoh’s algorithm (Gotoh, 1982) with space complexity of 𝒪 (*N* ^2^). For linear gap penalties, the algorithm is a variant of Needleman and Wunsch’s algorithm (Needleman and Wunsch, 1970) and requires a space complexity of *𝒪* (*N* ^2^). Both algorithms require additional *𝒪* (*N*) space to store the flags for insertions and deletions.

#### Preliminaries

The method traverses the guide tree in post-order and infers an alignment *A* for each node of the guide tree *τ*. The possible character states for each alignment site *k* = 1, …, |*A*| are recorded in *𝒫*_*k*_, denoted as parsimony set in the following. Additionally, we memorise deletions or possible insertions in two vectors *d, i* for each node, with |*d*|, |*i*| = |*A*|. The vector *d* can take the values 0 for no deletion or possible insertion, 1 for gap opening and 2 for gap extension.

Let *v* be an internal node of *τ*. Suppose the two child nodes *x* and *y* of *v* are leaves. In that case, the two sub-alignments *A*^*x*^ and *A*^*y*^ are given by the respective associated sequence. The parsimony sets 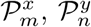 of the respective alignment sites *m* = 1, …, |*A*^*x*^|, *n* = 1, …, |*A*^*y*^| contain the character present at the particular site. We do not reconstruct evolutionary events at the leaf nodes, i.e., we set every entry of the vectors *d*^*x*^, *i*^*x*^ and *d*^*y*^, *i*^*y*^ to zero.

Since the affine gap penalty function is only subadditive, the optimal alignment between *A*^*x*^ and *A*^*y*^ up to column *k* is not necessarily in the optimal alignment up to column *k* + 1. Hence, we need to keep track of the optimal cost of all three prefix alignments in three separate scoring matrices with dimension (|*A*^*x*^| + 1) *×* (|*A*^*y*^| + 1). The optimal parsimony score for prefix alignment ending with a match between alignment columns is stored in *S*^*M*^. For prefixes ending with a column of *A*^*x*^ or *A*^*y*^ matched with a gap column, we store the cost in *S*^*X*^ and *S*^*Y*^, respectively. Additionally, we require three trace-back matrices per node, storing the optimal alignment path, each with dimension (|*A*^*x*^| + 1) *×* (|*A*^*y*^| + 1), denoted as *T* ^*M*^, *T* ^*X*^ and *T* ^*Y*^.

For each internal node *v*, the alignment *A*, the parsimony sets 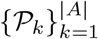 and the vectors *d, i* are found by the following alignment recursion:

#### Initialisation

We initialise the base cases for the alignments, that is, 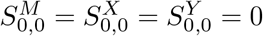 and

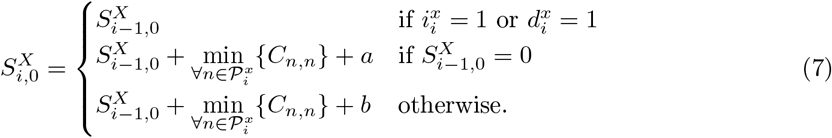

Analogous for 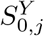.

The base cases that are not permitted or do not exist are initialised as follows: 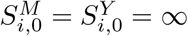 for all *i* ∈ {1, …, |*A*^*x*^|} and 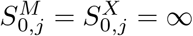 for all *j* ∈ {1, …, |*A*^*y*^|}.

#### Forward Phase

In the forward phase, we fill *S*^*M*^, *S*^*X*^, and *S*^*Y*^ for *i* = 1, …, |*A*^*x*^| and *j* = 1, …, |*A*^*y*^| according to the minimum parsimony score of either aligning column *i* of alignment *A*^*x*^ with column *j* of alignment *A*^*y*^ or aligning either one of the columns with a gap column. Columns flagged as insertions in the child node are always matched with a gap column since they cannot be descendants of ancestral characters. Consequently, we skip these columns in all three matrices to prevent the over-penalisation of insertion sites.

##### 1. Base case when neither 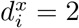 nor 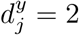

We can compute the score 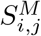 by aligning two columns *i* and *j* over the parsimony sets and the cost matrix by the following formula:

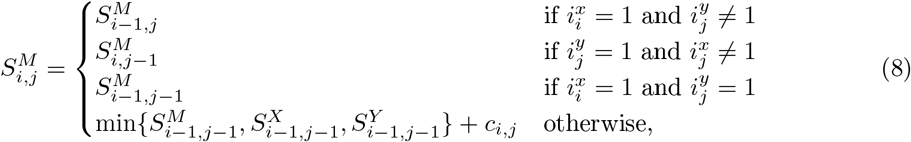

where

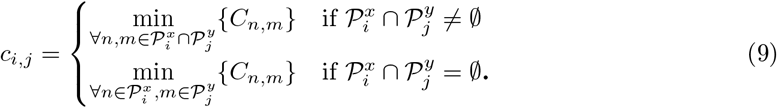

We record the optimal move inside the trace-back matrix *T* ^*M*^ according to the following formula:

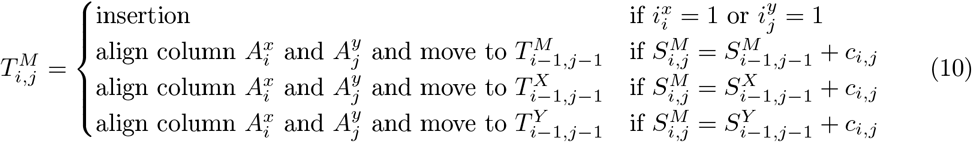

To calculate the score 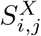 for aligning a column 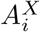 with a gap column, we have to follow the following steps. If the alignment site *i* is marked as a deletion in the child node *x*, i.e., 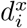 aligning 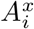 with a gap column is not penalised. Otherwise, aligning with a gap column is penalised with the gap opening penalty if the prefix alignment ends with a gap in the other sequence or with a match and the gap extension penalty if the prefix alignment ends with a gap in the same sequence. We calculate the score 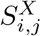 of aligning a column *i* of alignment *A*^*x*^ with a gap column as follows:

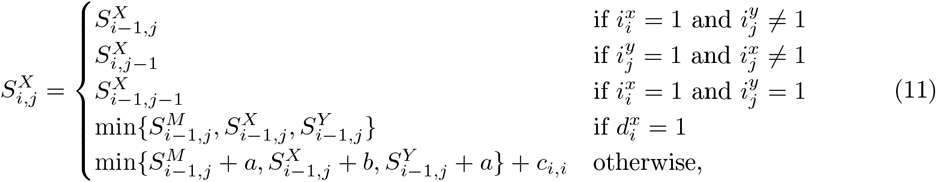

where

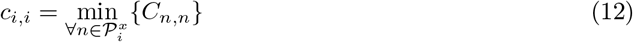

We store the optimal path in the trace-back matrix *T* ^*X*^ .

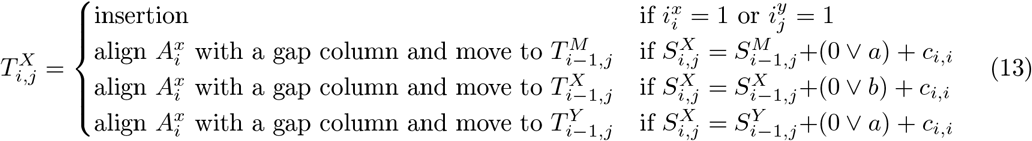

Analogous for *S*^*Y*^ and *T* ^*Y*^.

If the minimum value is not unique, we choose uniformly at random among the possibilities.

##### 2. Special case when 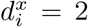 and or 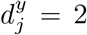

We are in a gap extension in either one or both of the child nodes, a match in the current node could break up a long gap in one of the child nodes. If this is the case, we must adjust the score by subtracting the gap extension penalty, which was added in the previous step and adding a gap opening penalty. In other words, we must check if a long deletion event is separated into an insertion and deletion event and adjust the score accordingly. To do this, we have to trace back through the current steps skipping insertion sites and find the previous residue that could belong to the same evolutionary event.

First, we compute the score 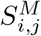 for aligning two columns *i, j*. Let us assume that only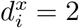.

If we consider the prefix alignment that matches a column in *x* with a gap column, i.e., 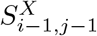, we always have to adjust the score. Since the previous column was either a gap extension or opening, matching the current column with a column containing a residue means we break up the long event in the child node. The adjusted score is

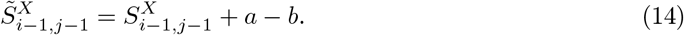

If we consider the prefix alignment where a residue column in *y* was matched with a gap column, we need to trace back to find either the pointer to the matrix *S*_*X*_ or *S*_*M*_. We only need to adjust if it does not point to *S*_*M*_ because the event would be separated into a previous insertion and deletion at the current site. Let 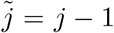 and 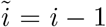, while we stay in *T* ^*Y*^ we set 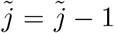 if 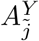 is marked as an insertion or if it is matched with a gap column, and we set 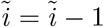 if 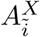 is marked as an insertion. The adjusted score is

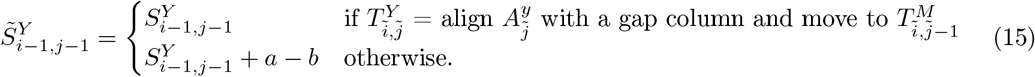

In summary, we get,

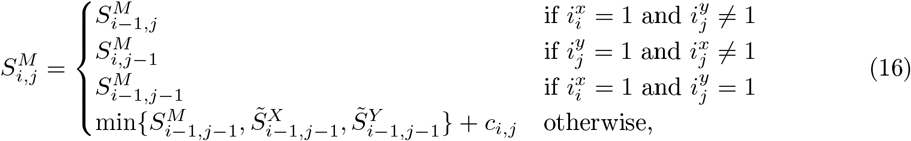

where

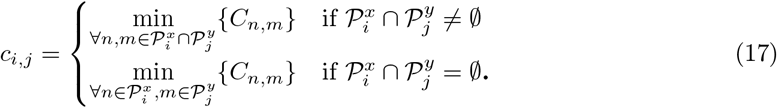

Analogously for 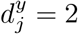. If both residues are in an open gap in the child node, i.e., 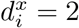 and 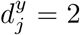, we must trace back through both matrices *T* ^*Y*^ and *T* ^*X*^. Since we close two open gaps in the current node, the adjusted score is given by

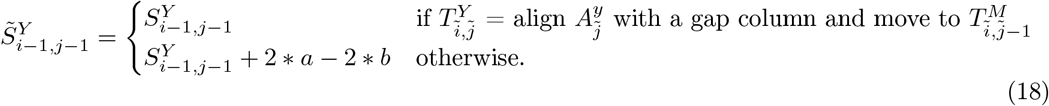

for the trace back through *T* ^*Y*^. Analogous for *T* ^*X*^. The trace-back matrices are found analogous to the base case.

Second, we compute the score 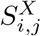 Let us again assume that we are in an open gap in 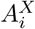, i.e. 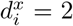. Let’s consider the prefix alignment that ends with a match of both columns. We always have to adjust the score since we separate the long event into a deletion in the previous column and an insertion in the current column. The adjusted score is

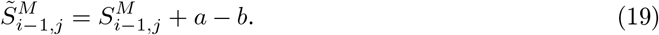

For the prefix alignment that ends with matching the column 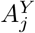with a gap column, we have to trace back and only adjust the score if we do not point to *T* ^*X*^. Let 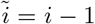 and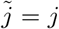, while we stay in *T* ^*Y*^, we set 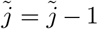 if 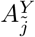 is marked as an insertion or if it is matched with a gap column, and we set 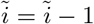 if 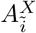 is marked as an insertion. The adjusted score is

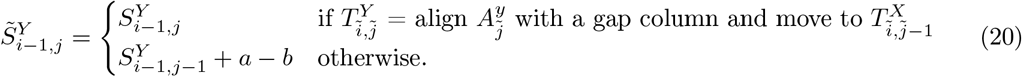

In summary, we get

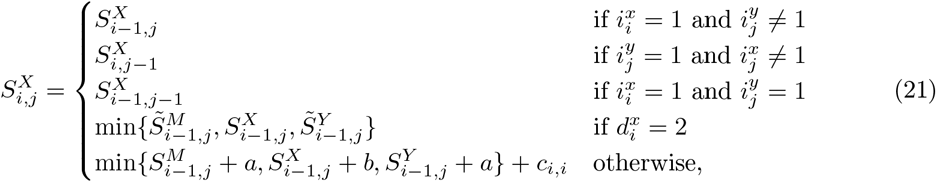

where

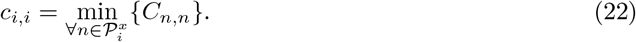

Analogous for 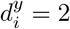 and 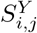. The trace-back matrix is found analogous to the base case.

#### Backward Phase

We trace back through the matrices *T* ^*M*^, *T* ^*X*^, *T* ^*Y*^ for the backwards phase. The starting matrix is chosen based on 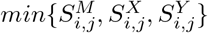, with *i* = |*A*^*x*^|, *j* = |*A*^*y*^|. We find the optimal alignment *A* starting from the last alignment column.

If the step is labelled as an insertion in any trace-back matrix and both columns are marked as an insertion in the child nodes, we form two new alignment columns. We align 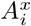 and 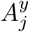 with a gap column, let the respective new parsimony set only include the gap character and flag the resulting columns as insertion sites. Subsequently, we decrement *i* and *j* and proceed with the next trace-back step without switching matrices. If only one column is labelled as an insertion, we align the respective column with a gap column. The resulting parsimony set per column only includes the gap character, and the column is flagged as an insertion. Here only the index of the aligned column is decremented, and we stay in the same trace-back matrix.

If the current matrix is *T* ^*M*^, we align the two columns 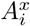 and 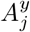. The parsimony set of possible character states is either given by the intersection or union of the two parsimony sets of the two columns. Unions correspond to substitution events, while intersections represent a match. Since we either match identical characters or substitute one character for another, the site is neither labelled as insertion nor deletion in the resulting alignment. Consequently, we decrement *i* and *j* and switch the trace-back matrix depending on the optimal prefix alignment.

If the current matrix is *T* ^*X*^, we align the column 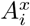 with a gap column. If 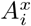 was not labelled as deletion, i.e., 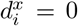, we determine the parsimony set for the new alignment column by 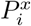 and label the resulting column as deletion. Otherwise, the column was flagged as a deletion in the child node and aligned with a gap column without penalty in the current node. Therefore, the optimal solution is to relabel the column in the resulting alignment as insertion. The parsimony set includes the gap character since inserted sites do not have homologous ancestor sites. In subsequent alignment steps, we label the column as an insertion to prevent alignment with non-homologous characters. Since we only used a column of alignment *A*^*x*^, we decrement *i* and switch between trace-back matrices if necessary. Analogous for *T* ^*Y*^.

We summarise the calculation of parsimony sets and the determination of insertion and deletion flags in the following formula:

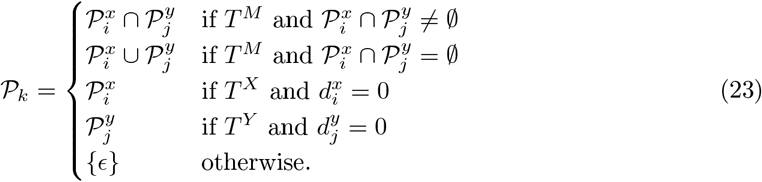

and

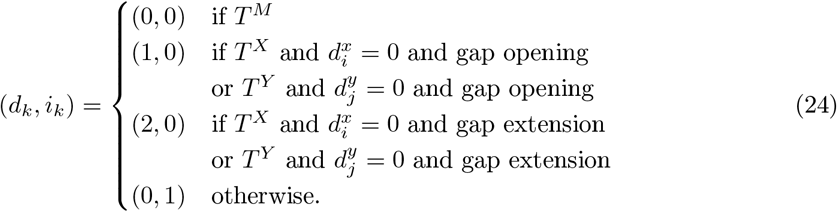

### 5.3 Cost matrix and gap penalties

Both methods are compatible with transition probability matrices *P* (*t*) of nucleotide or amino acid substitution models. The time *t* is a measure of the distance *d* between the two sequences. The cost *C*_*i,j*_ for a match between character *i* and character *j* is given by (Yang, 2006):

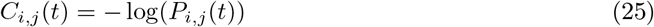

To save computational time, the cost matrix is not recalculated for each branch length. Instead, one cost matrix for an average distance is used, or four cost matrices are calculated based on the branch lengths’ 20th, 40th, 60th and 80th percentile. The cost matrix derived from the distance closest to the length of the branch is chosen during inference. Choosing suitable gap penalties posed quite a challenge. We refrained from tuning parameters to one specific dataset to ensure consistency and comparability between the experiments. Instead, we used the average substitution cost multiplied by 2.5 for the gap opening and the 0.5 for the gap extension penalty based on recommendations by Altschul et al. (Altschul and Erickson, 1986).

### 5.4 Code

The code is available from https://github.com/acg-team/indelMaP.

### 5.5 Simulated sequence data with indels

To validate and compare our methods to state-of-the-art methods, we followed a previous simulation study that assessed the impact of alignment accuracy on ASR accuracy (Vialle et al., 2018). We used INDELible v1.03 (Fletcher and Yang, 2009) to simulate 54 scenarios using a combination of realistic evolutionary parameters summarised in Table 2.

**Table 2:** The parameters used for simulating random trees (left) and associated amino acid sequences (right) with INDELible. Vertical lines indicate parameter values which were combined to create 24 scenarios.

|  |  |  |  |
| --- | --- | --- | --- |
| **Number of taxa** | 16 \| 32 \| 64 | **Substitution model** | WAG |
| **Birth rate** | 6 | **Root sequence length** | 408 |
| **Death rate** | 3 | **Indel model** | Zipfian distribution, factor 1.7 |
| **Sampling fraction** | 0.01 \| 0.25 \| 0.99 | **Max. indel length** | 20 |
| **Mutation rate** | 0.8 \| 1.2 \| 1.7 | **Indel rate** | 0.01 \| 0.05 |

First, we simulated ten random trees for each of the 27 tree parameter combinations, resulting in 270 trees. The random trees are simulated following a birth-death process for 16, 32 or 64 taxa. The tree height varies over the mutation rate, representing the average number of substitutions per site. The data used for the benchmark can be found at https://github.com/acg-team/indelMaP/benchmark.

Subsequently, we simulated ten sets of extant sequences and their evolutionary history for each of the ten random trees and two indel rates, resulting in 5400 sequence sets. The indel rate represents insertion and deletion rates, which are equal and relative to an average substitution rate of one. Substitutions are modelled under the WAG model (Whelan and Goldman, 2001), and indels according to the power law with a constant factor of 1.7 and a maximum indel length of 20.

### 5.6 Phylogenetic inference tools

We compared our methods with current state-of-the-art inference tools. For inferring an ASR estimate, we used FastML v3.11 (Ashkenazy et al., 2012) without branch length optimisation and with likelihood-based indel reconstruction, GRASP (Foley et al., 2019), ARPIP (Jowkar et al., 2022) and Historian (Holmes, 2017) with the option -band 0. Additionally, we compared results to a conventional parsimony score treating gaps as an additional state. For MSA estimation, we used the conventional aligner MAFFT v7.490 (Katoh, 2002) run with the –auto option, PRANK v.170427 (Löytynoja, 2014) with the option +F, ProPIP (Maiolo et al., 2021) without optimisation and Historian with default parameters. IndelMaP was run with default parameters for ASR and MSA estimation. All methods were only evaluated when given the true tree for their estimation. If possible, the substitution model was given to the method, except for MAFFT, which uses BLOSUM62 as a default scoring matrix.

### 5.7 Quality measures

To assess the quality of the ASR estimate, we chose to adopt an accuracy measure previously used in (Paten et al., 2008; Vialle et al., 2018). The reconstructed sequences are compared to the true sequences to calculate an overall accuracy measure. Pairwise alignment is only necessary if the reconstructed and true ancestral sequences differ in length. This is sometimes the case for Historian since the tool uses the provided MSA only as a guide for ASR estimation. To keep the inferred gap pattern, we treat gaps as an additional state during pairwise alignment and introduce an additional gap character. Sequences are aligned using a Needleman and Wunsch algorithm (Needleman and Wunsch, 1970).

The overall accuracy is calculated based on three error scores:

i. Substitution error: the sum of mismatched residues.
ii. Insertion error: the sum of residues reconstructed in the inferred sequence aligned with a gap character in the true sequence.
iii. Deletion error: the sum of gap characters in the reconstructed sequence aligned with a residue in the true sequence.
iv. Alignment error: the sum of residues aligned to the additional gap character in either the true or reconstructed sequence.

All error scores, except the insertion error, are scaled by the length of the true sequence (without gap characters). The insertion error is scaled by the number of gap characters in the true sequence. The overall accuracy is calculated by deducting the sum of all errors from one.

As a means of evaluating the accuracy of MSA estimation, we employed the Sum-of-Pairs (SPS) and Total Column Score (TCS) measures, also known as Column Score (Thompson et al., 1999). The SPS calculates the proportion of correctly aligned residue pairs, while the TCS gives the proportion of accurately aligned columns for the entire alignment. We used qscore v2.1 (available for download at http://www.drive5.com/qscore) to calculate both scores.

To determine the percentage of accurately inferred insertion and deletion events, we compared the estimated events (i.e. location on the tree, length, and aligned residues) to the simulated ground truth.

All quality measures computed for a given estimated guide tree only apply to subtrees that exist in both the simulated and estimated tree.

### 5.8 Amino acid sequences of gp120

The data set consists of 761 gp120 sequences of subtype B of human immunodeficiency virus type 1 (HIV-1) from European countries downloaded from the Los Alamos National Laboratory HIV Sequence Database http://www.hiv.lanl.gov/, plus the reference sequence HXB2. We extracted only one sequence per patient and excluded sequences where the year or subtype was not indicated. We utilised the accession number to query the GenBank (Benson et al., 2012) and retrieve a more accurate sampling date if one was available. After obtaining the data set, we used a bioNJ guide tree to estimate an initial MSA with indelMaP. The cost matrix used for the analysis was derived from the HIVb model (Nickle et al., 2007), an amino acid substitution model for between-host analysis of HIV-1 sequences. The initial MSA was given to the phylogenetic tree inference tool, IQ-tree (Nguyen et al., 2015) to infer a dated phylogenetic tree (To et al., 2016). We used the option to outgroup root the tree with the reference sequence and the HIVb model. After inferring the tree, we used it as a guide tree for indelMaP, estimated an MSA and reconstructed the ancestral sequences and indel events with a HIVb cost matrix. The variable regions of the gp120 sequences were extracted based on the alignment to the reference sequence. For calculating the fraction of inserted or deleted sites per site per year and for indel lengths, branch lengths close to zero of the dated tree were excluded. The code used for the analysis can be found at https://github.com/acg-team/indelMaP/gp120-indel-pattern-analysis.

### 5.9 Data Availability Statement

The data underlying this article are available at https://github.com/acg-team/indelMaP.

## A Supplementary Material

### A.1 Ancestral sequence reconstruction accuracy for tree height 1.2 and 1.7

**Figure SM1:**
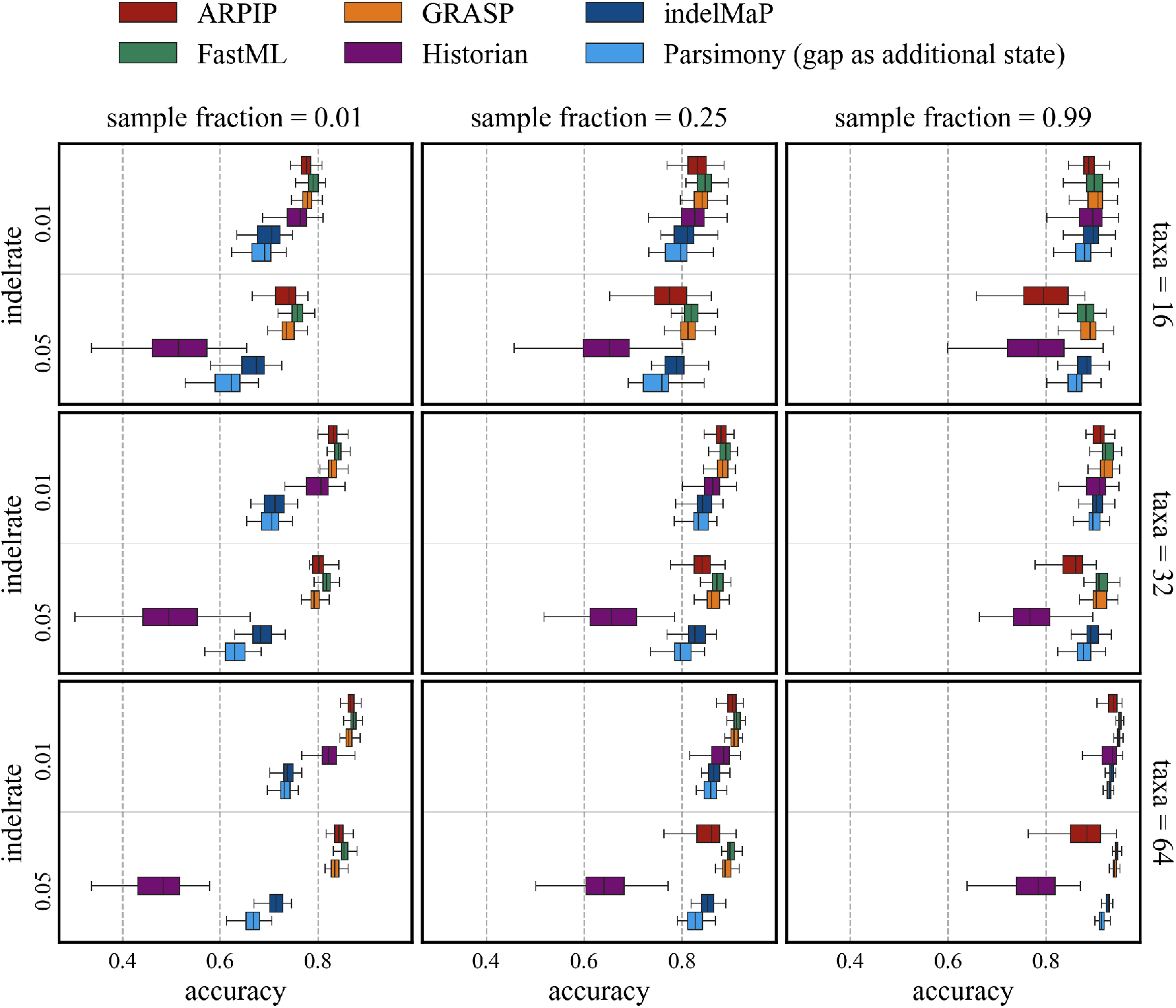
Overall character reconstruction accuracy for all parameter combinations for tree height 1.2.

**Figure SM2:**
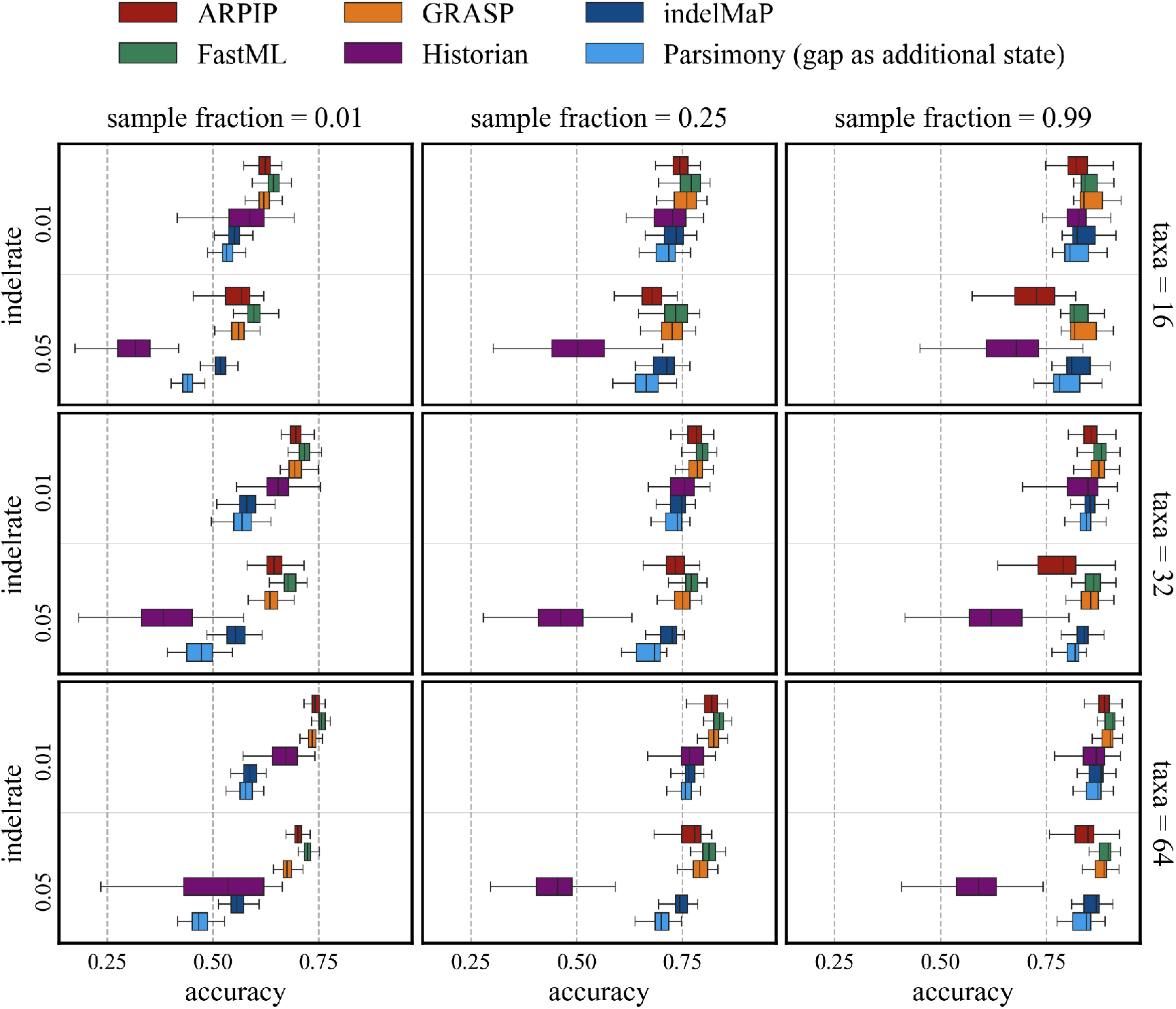
Overall character reconstruction accuracy for all parameter combinations for tree height 1.7.

### A.2 Ancestral sequence reconstruction substitution, insertion and deletion error for tree height 0.8

**Figure SM3:**
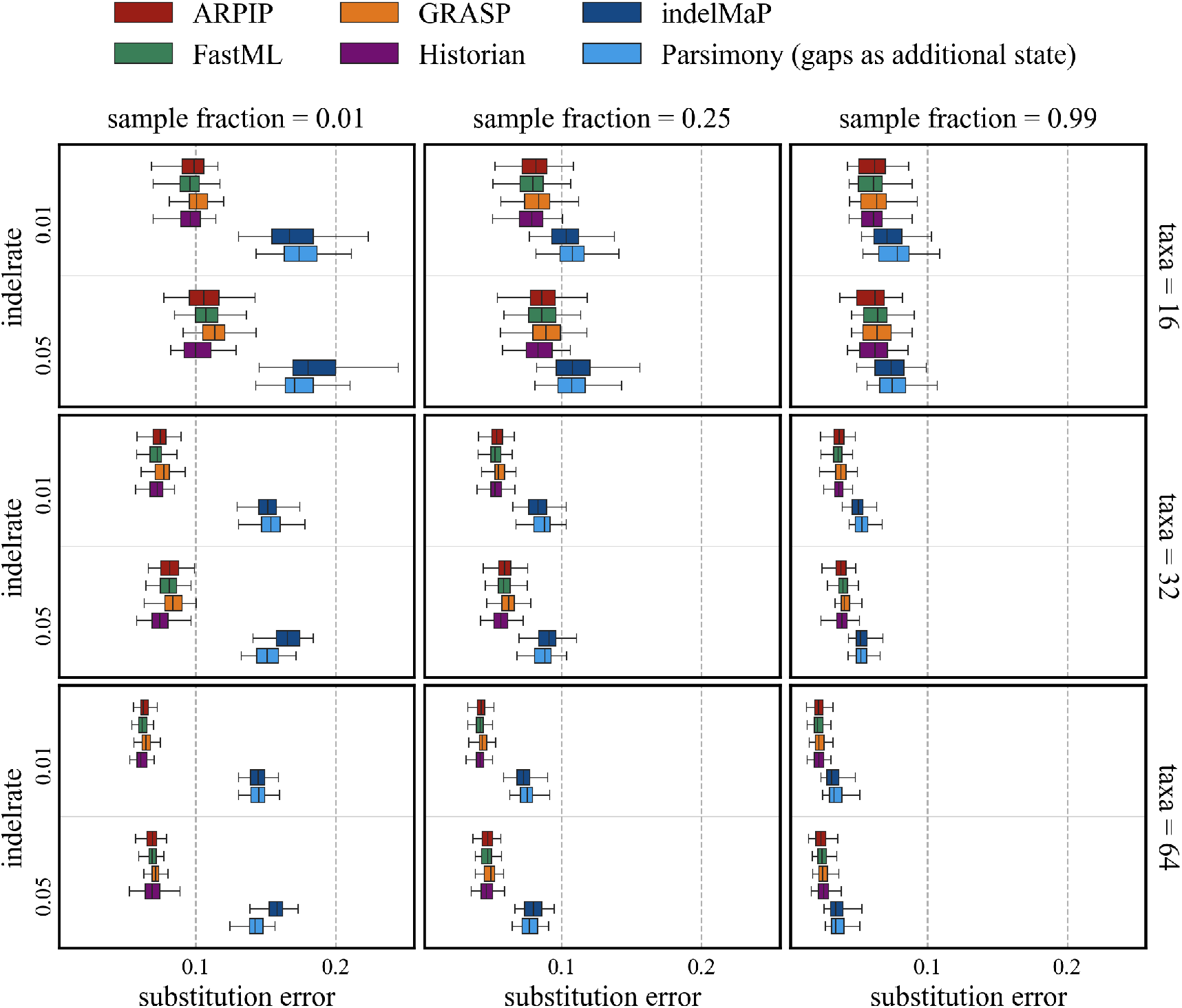
Substitution error for all parameter combinations for trees with tree height 0.8

**Figure SM4:**
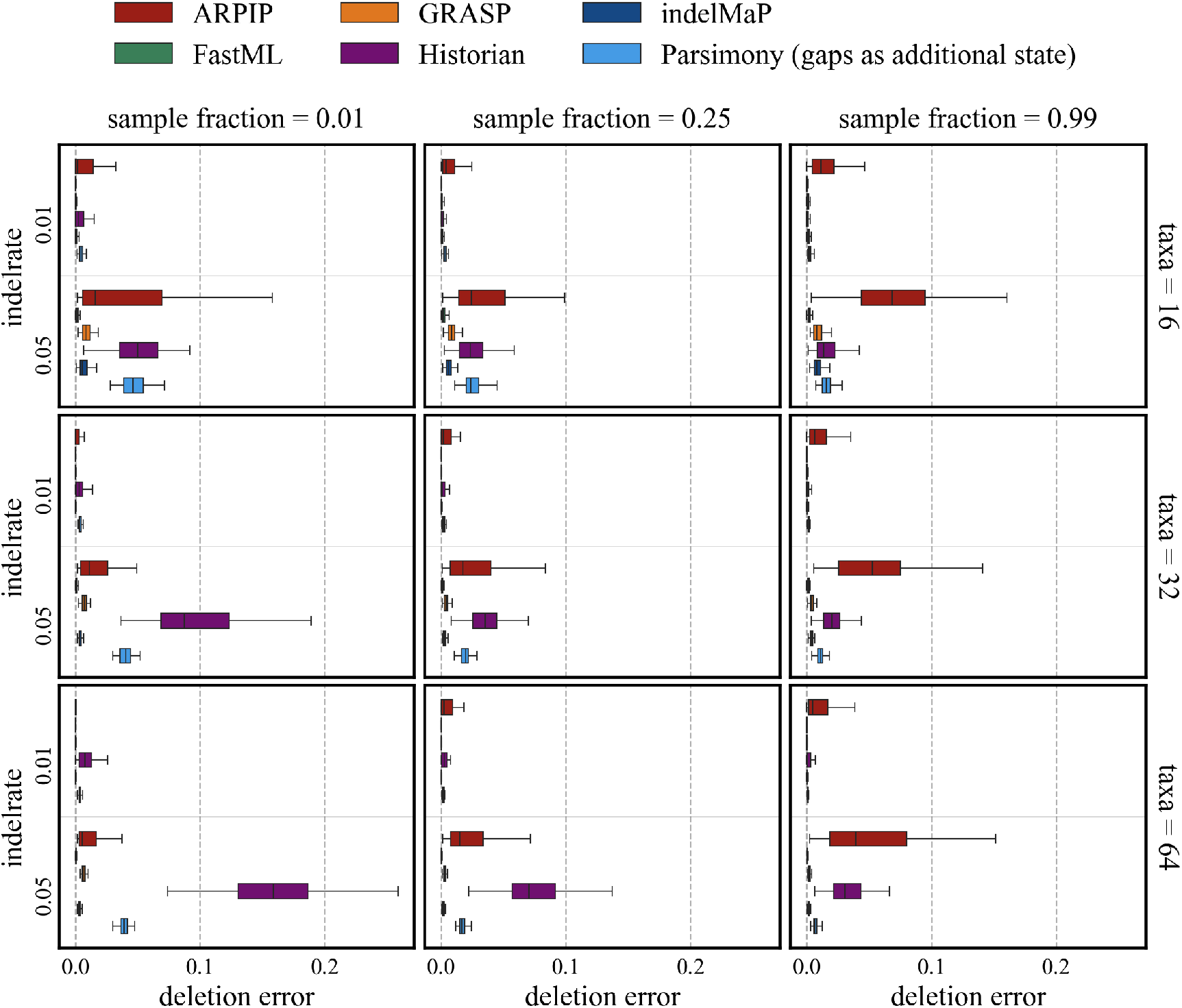
Deletion error for all parameter combinations for trees with tree height 0.8

**Figure SM5:**
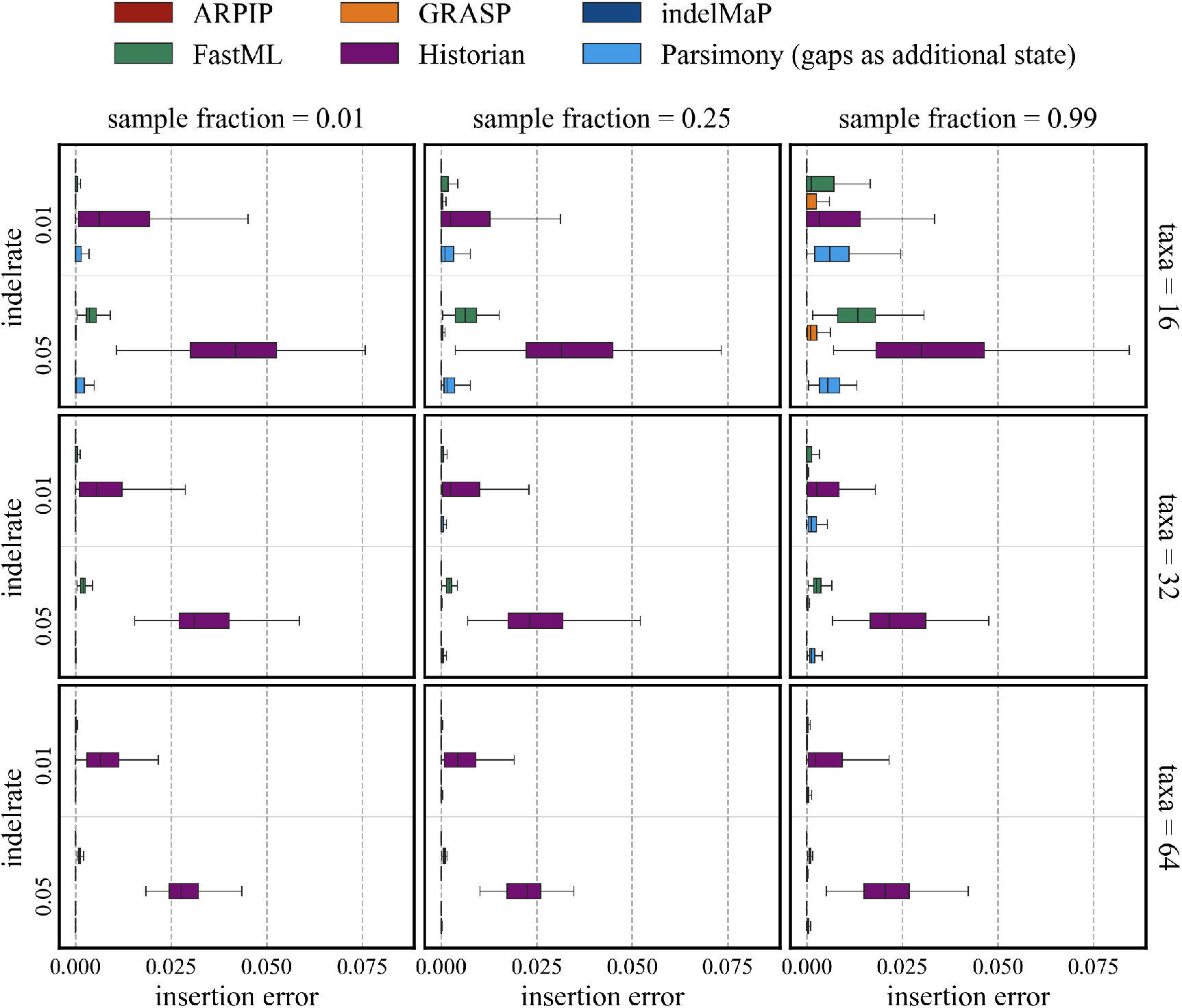
Insertion error for all parameter combinations for trees with tree height 0.8

### A.3 Ancestral sequence reconstruction substitution, insertion and deletion error for tree height 1.2

**Figure SM6:**
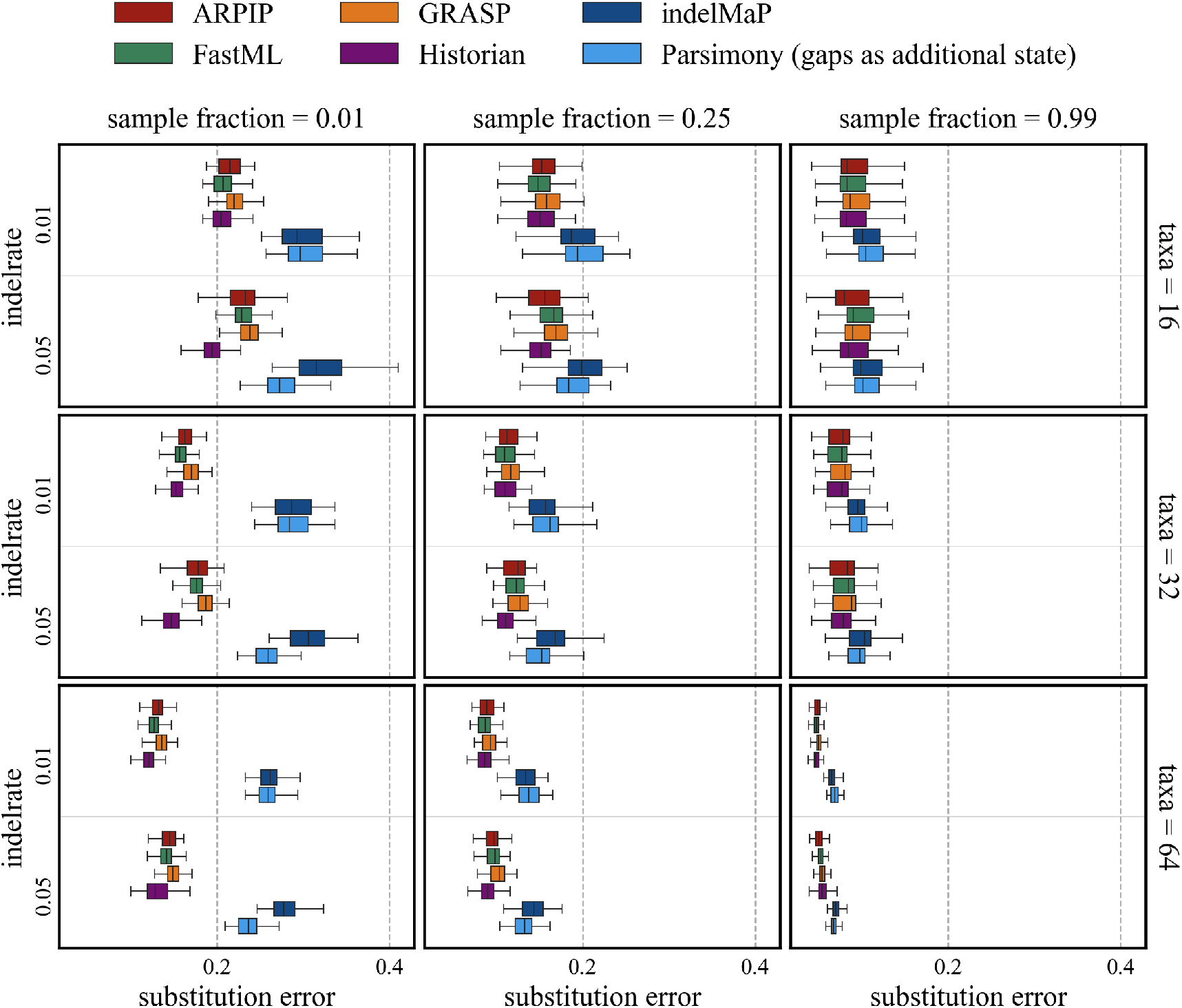
Substitution error for all parameter combinations for trees with tree height 1.2

**Figure SM7:**
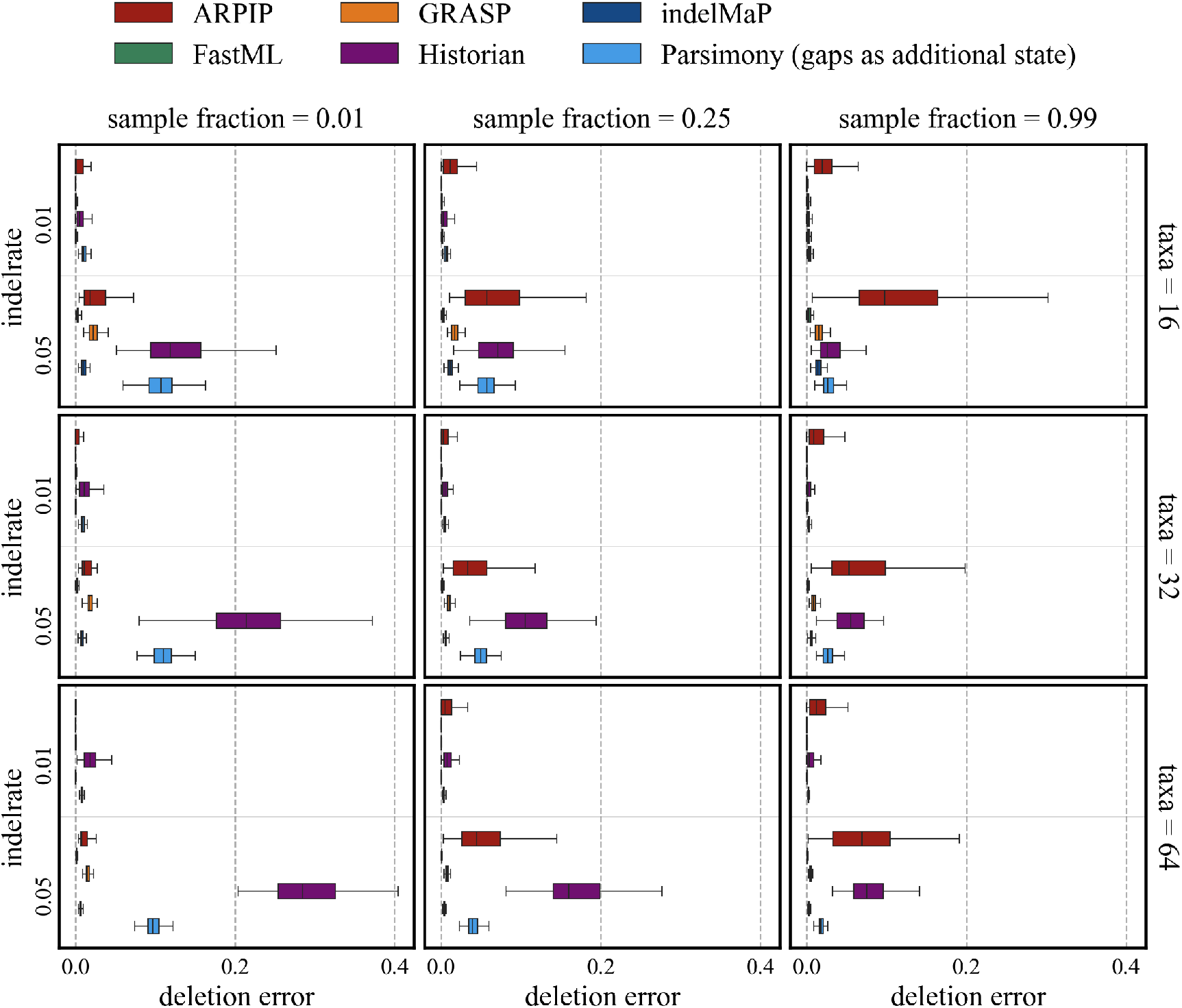
Deletion error for all parameter combinations for trees with tree height 1.2

**Figure SM8:**
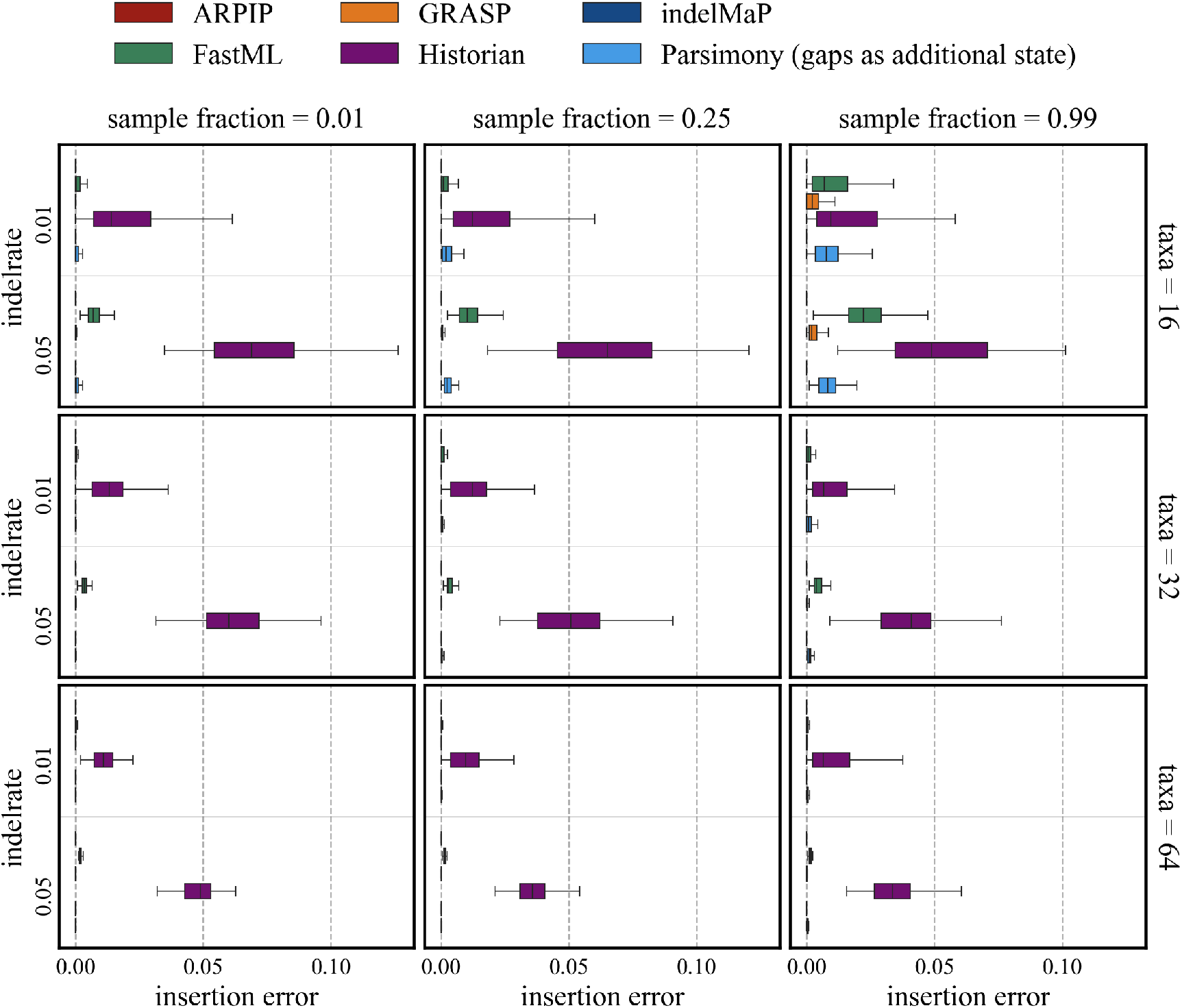
Insertion error for all parameter combinations for trees with tree height 1.2

### A.4 Ancestral sequence reconstruction substitution, insertion and deletion error for tree height 1.7

**Figure SM9:**
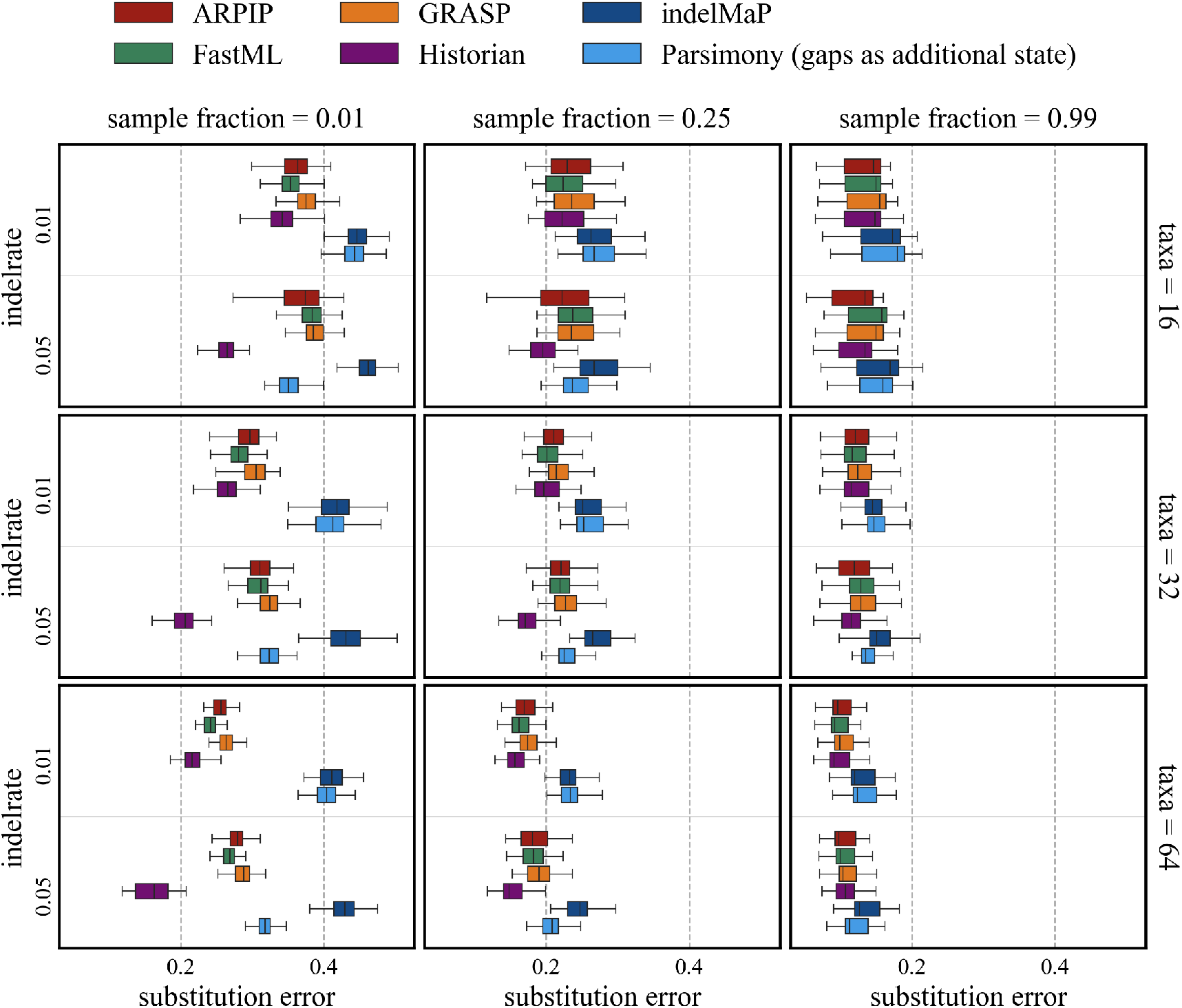
Substitution error for all parameter combinations for trees with tree height 1.7

**Figure SM10:**
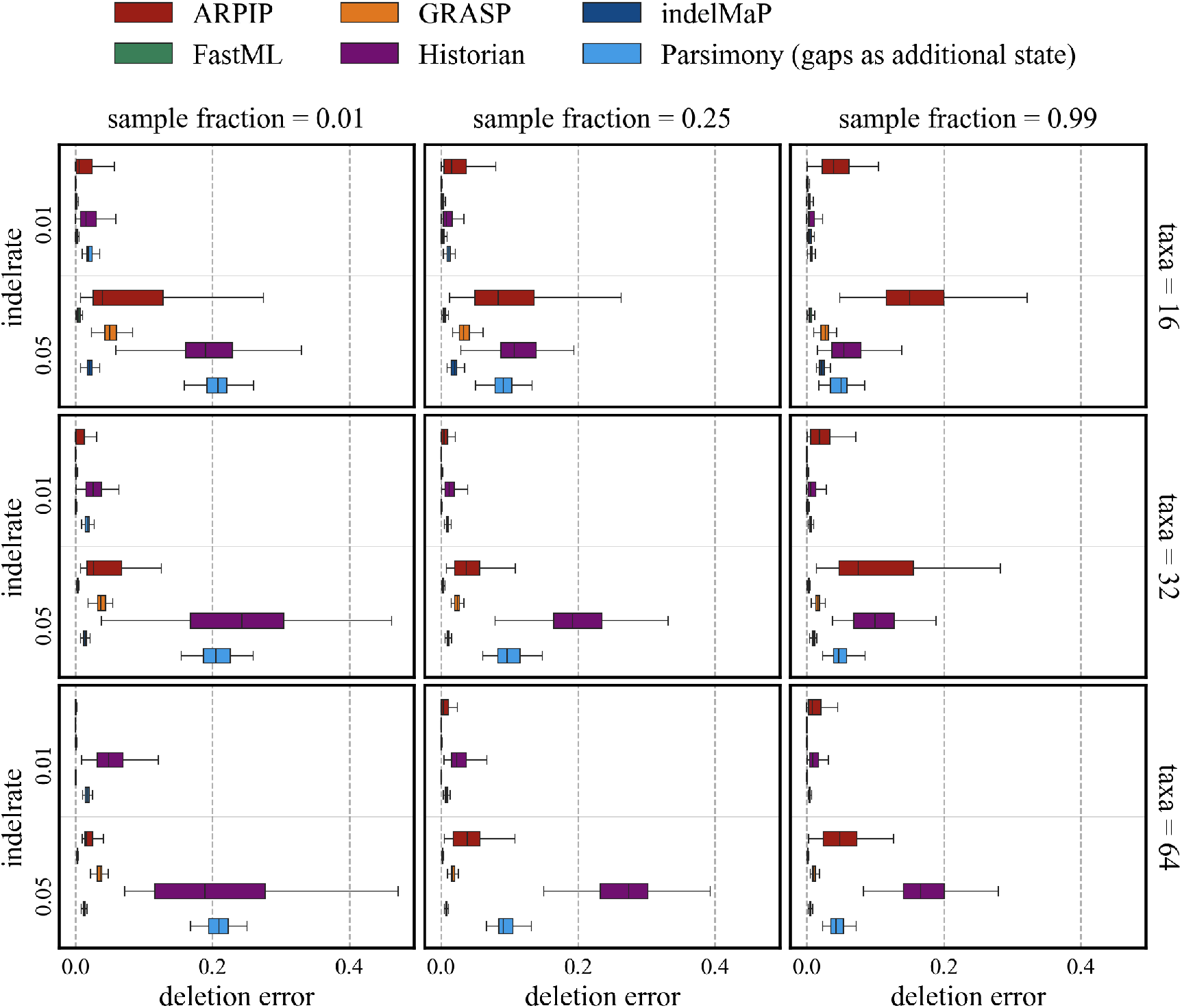
Deletion error for all parameter combinations for trees with tree height 1.7

**Figure SM11:**
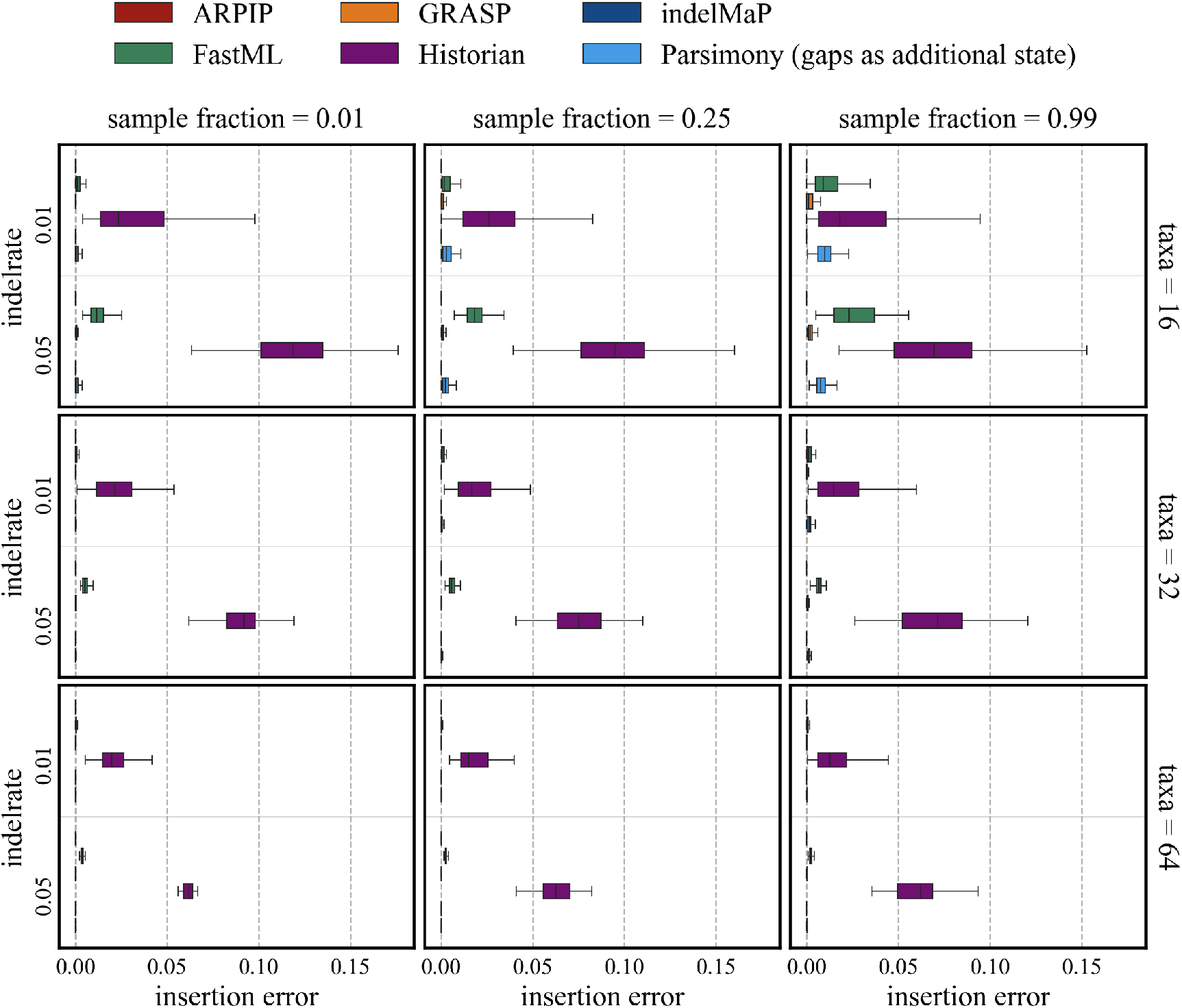
Insertion error for all parameter combinations for trees with tree height 1.7

### A.5 Correlation between error scores and distance to child nodes

**Figure SM12:**
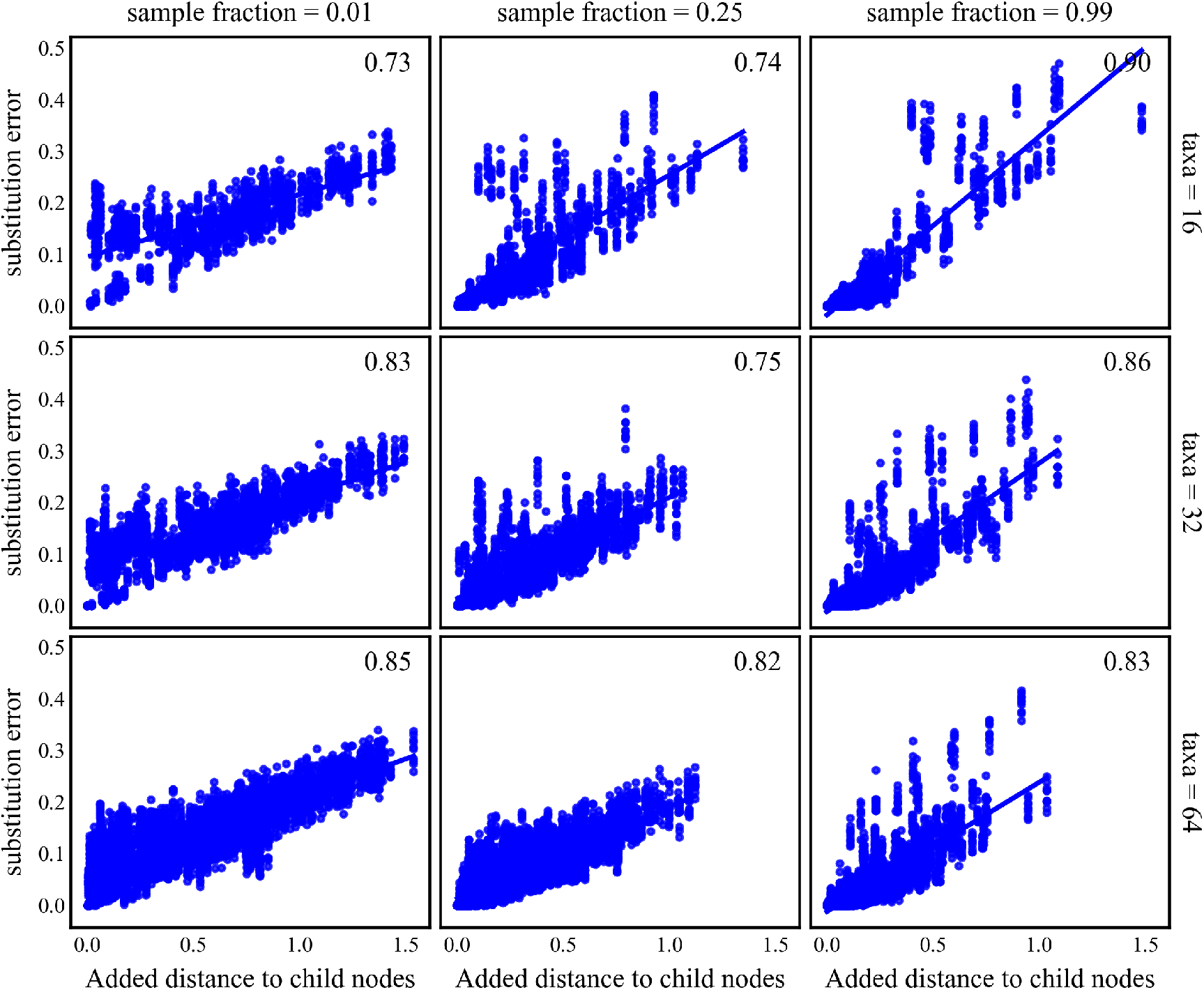
Correlation between added distance to the child nodes and substitution error for all parameter combination with tree height 0.8 and indel rate 0.01.

**Figure SM13:**
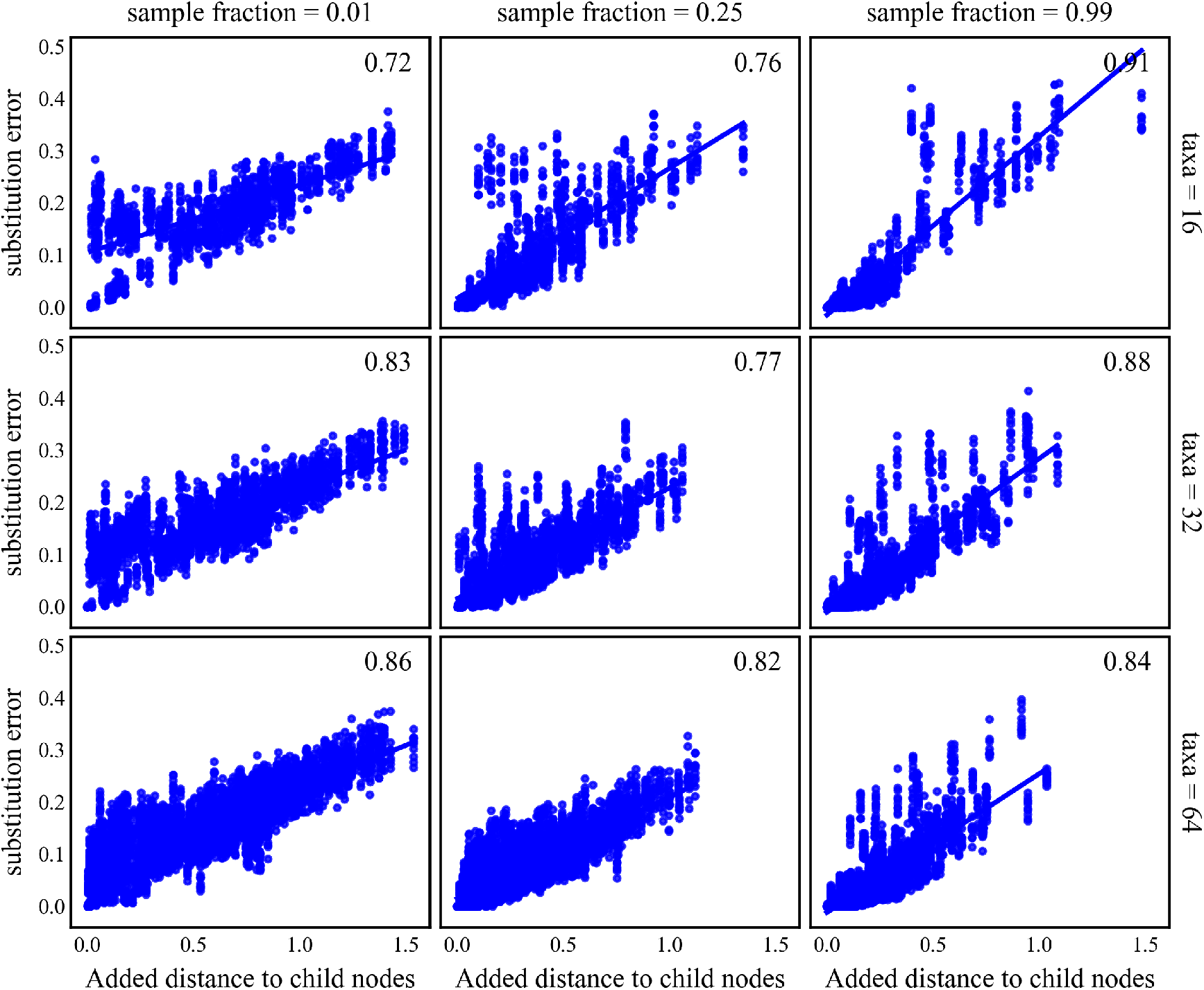
Correlation between added distance to the child nodes and substitution error for all parameter combination with tree height 0.8 and indel rate 0.05.

**Figure SM14:**
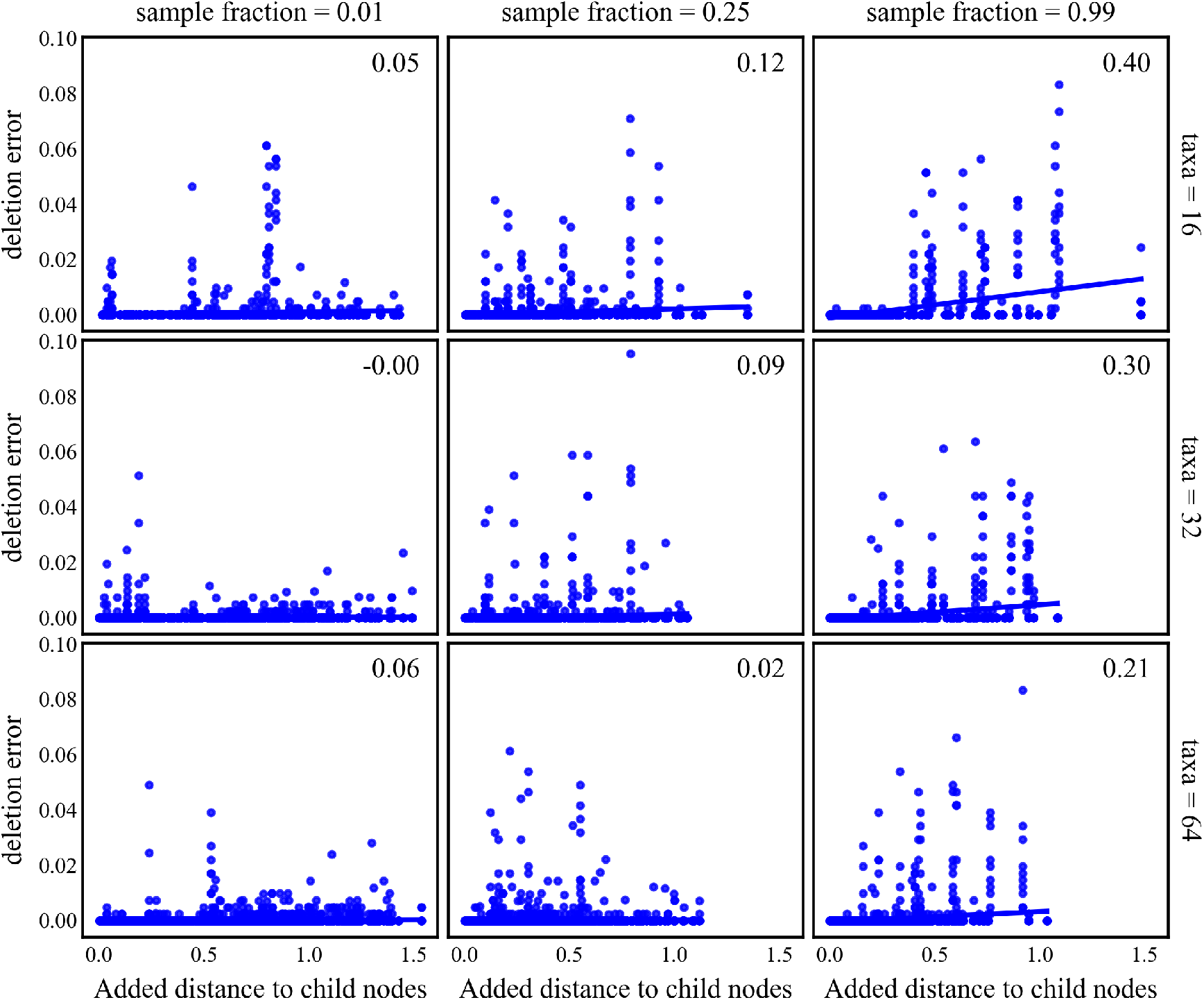
Correlation between added distance to the child nodes and deletion error for all parameter combination with tree height 0.8 and indel rate 0.01.

**Figure SM15:**
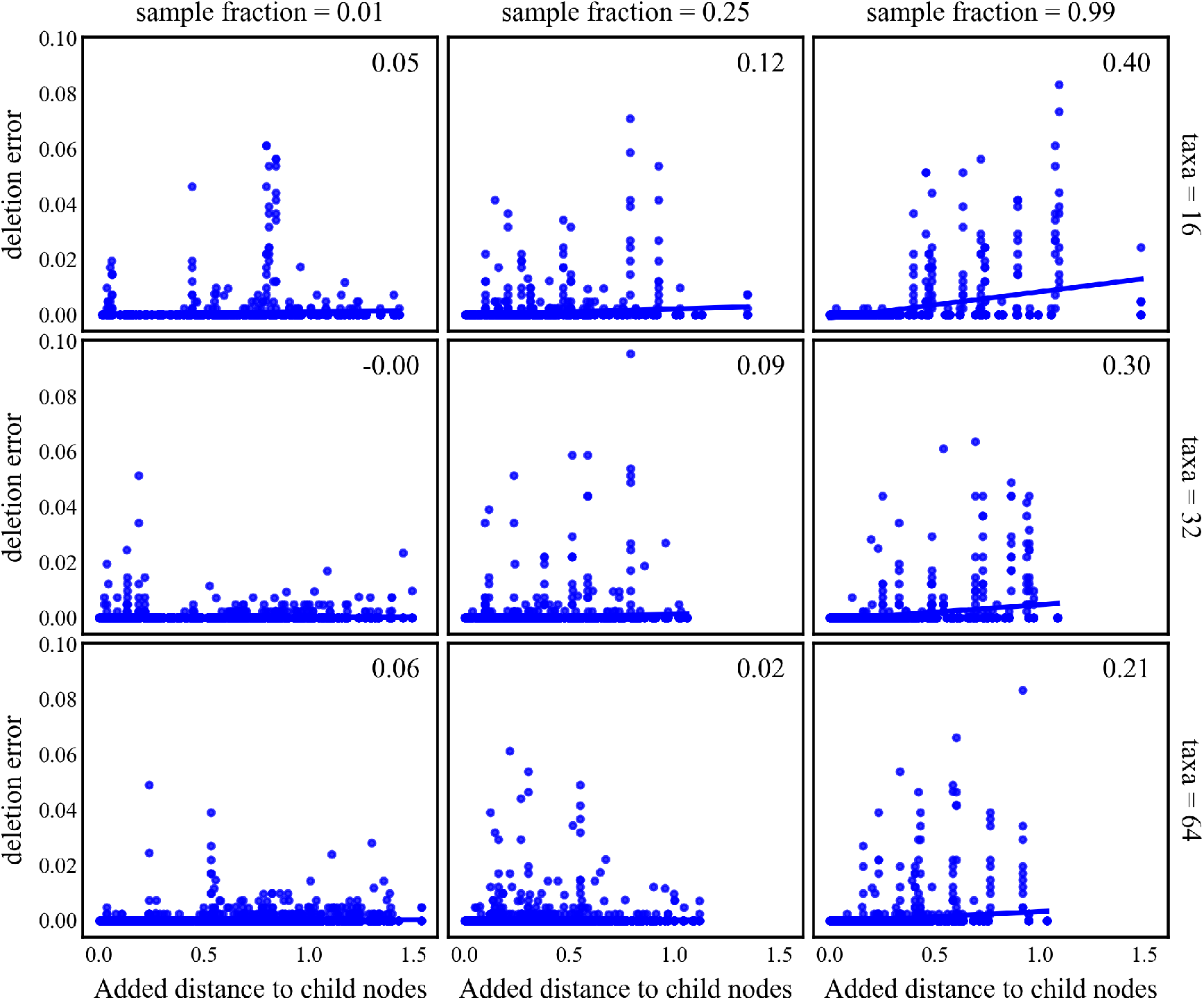
Correlation between added distance to the child nodes and deletion error for all parameter combination with tree height 0.8 and indel rate 0.05.

### A.6 Multiple sequence alignment quality for tree height 1.2 and 1.7

**Figure SM16:**
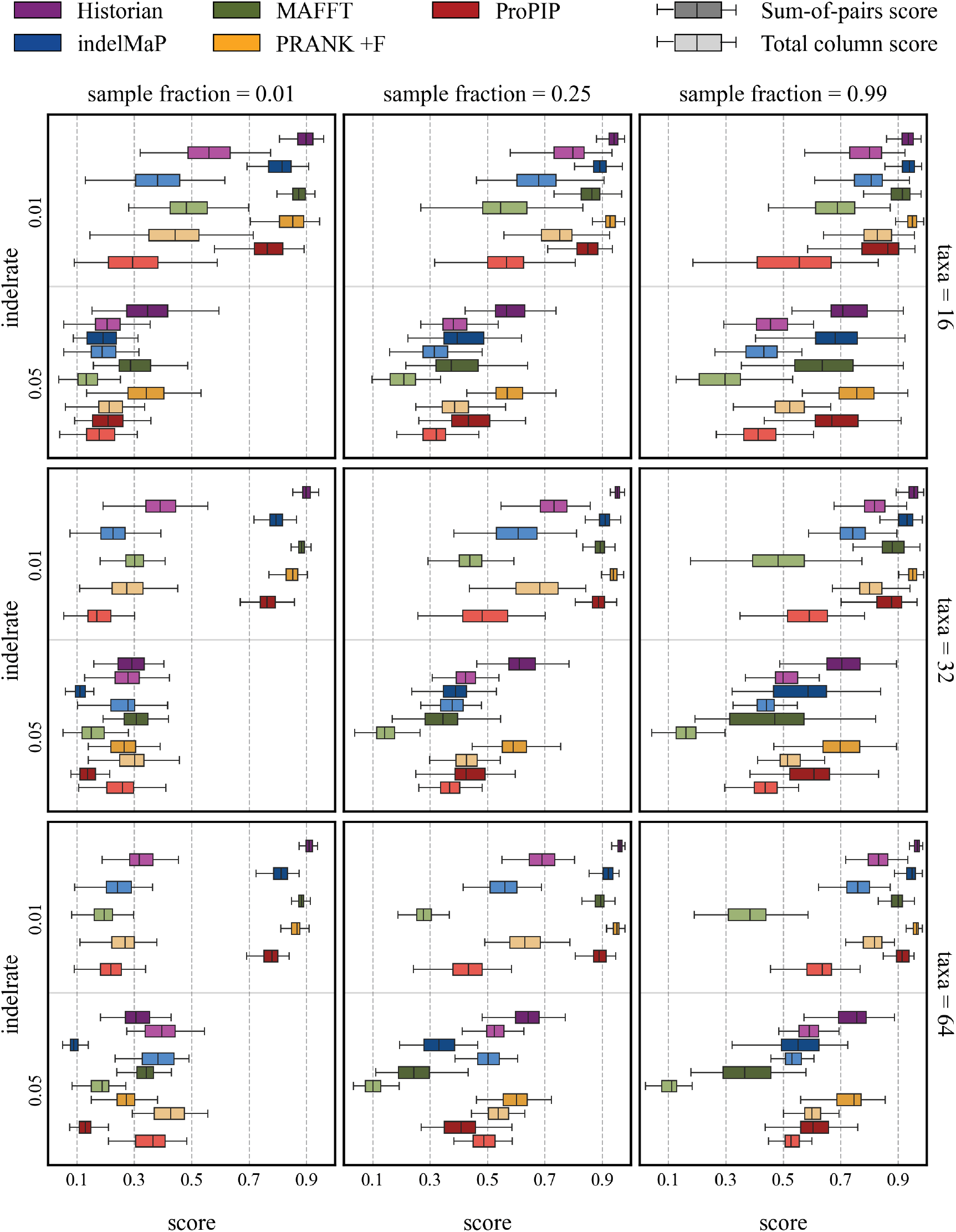
SPSs and TCSs scores for all parameter combinations with tree height 1.2.

**Figure SM17:**
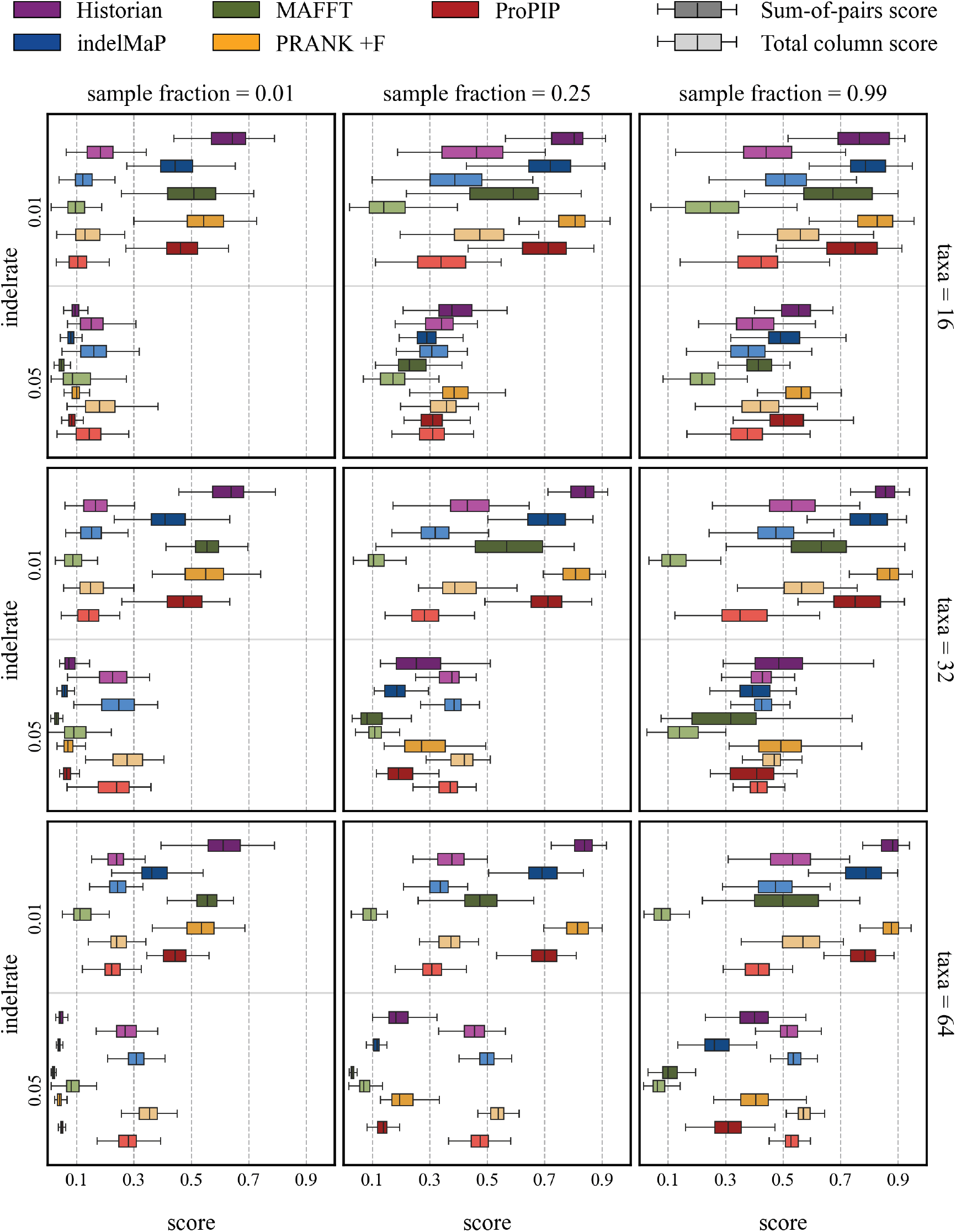
SPSs and TCSs scores for all parameter combinations with tree height 1.7.

**Figure SM18:**
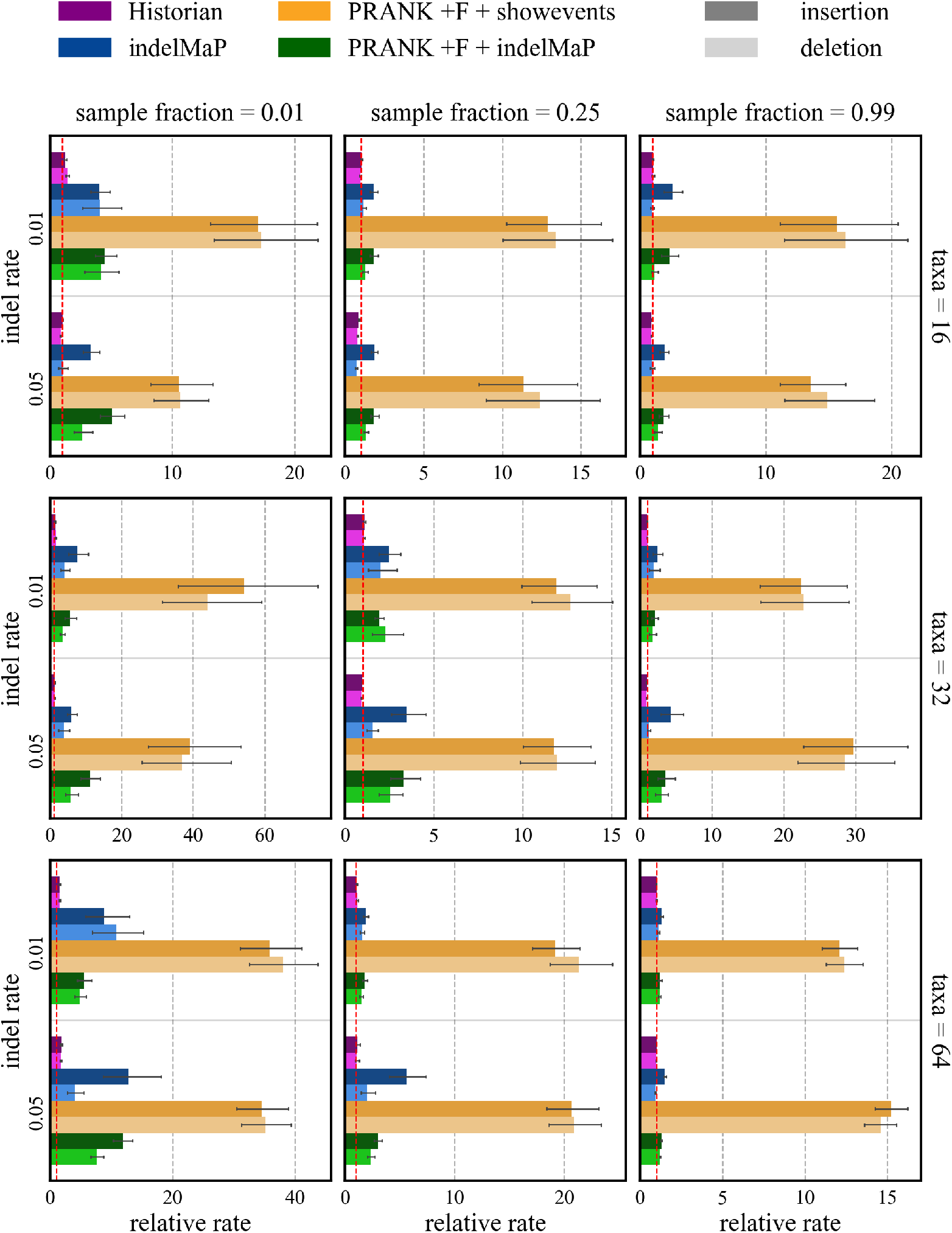
Relative insertion and deletion rates for all parameter combinations with tree height 1.2

**Figure SM19:**
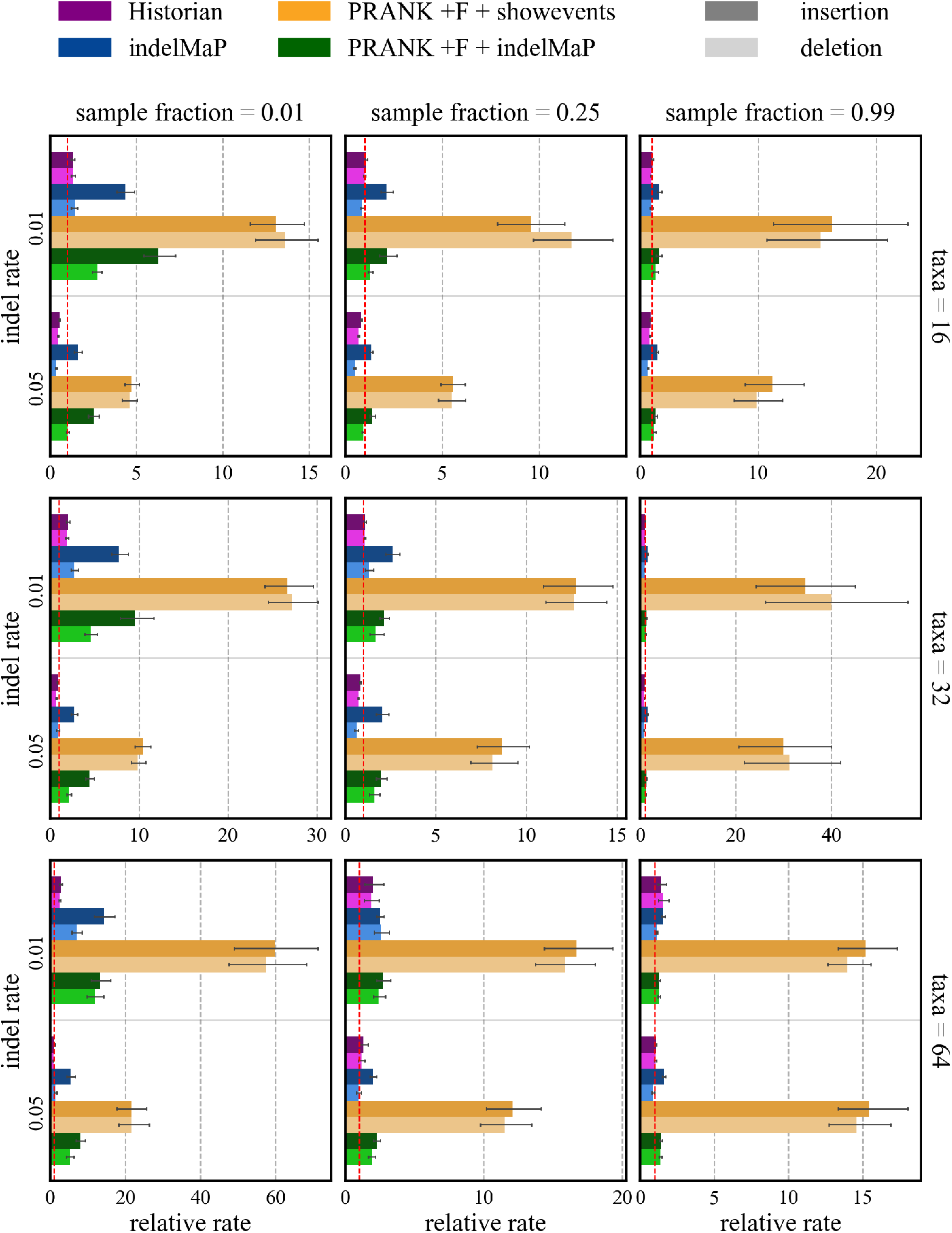
Relative insertion and deletion rates for all parameter combinations with tree height 1.7

**Figure SM20:**
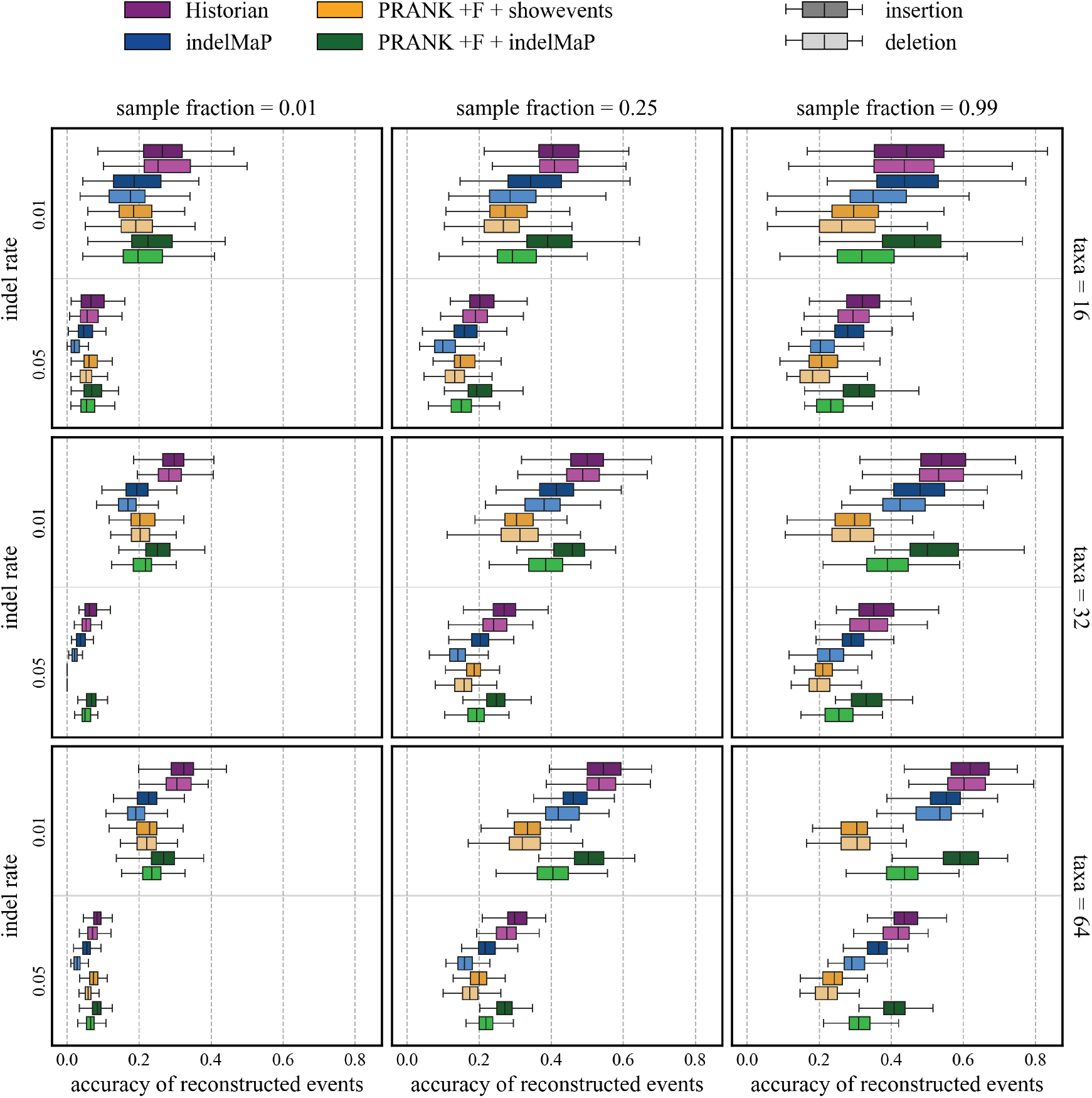
Proportion of accurate inferred insertion and deletion rates for all parameter combinations with tree height 1.2.

**Figure SM21:**
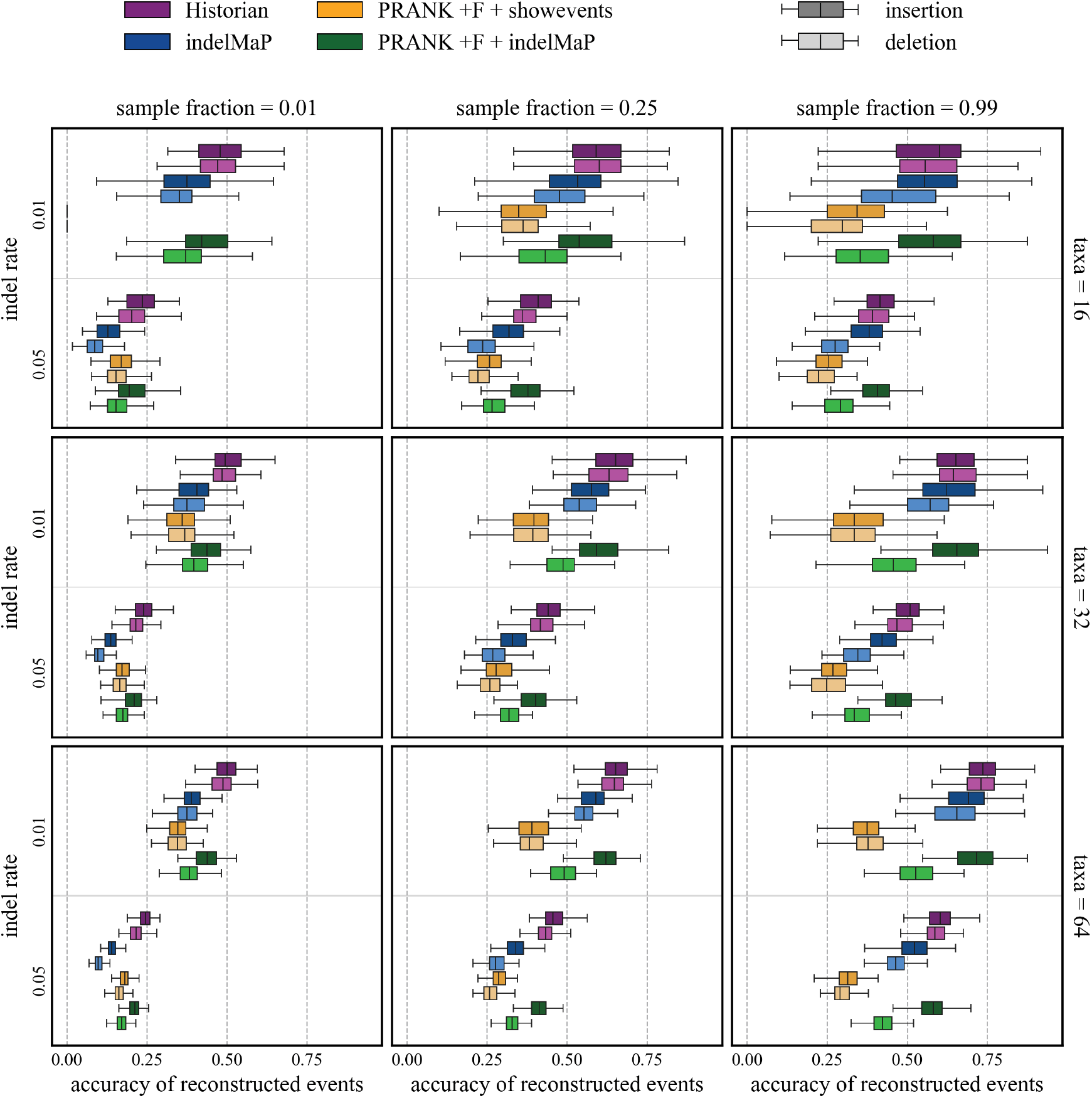
Proportion of accurate inferred insertion and deletion events for all parameter combinations with tree height 1.7.

### A.7 Time benchmark

**Figure SM22:**
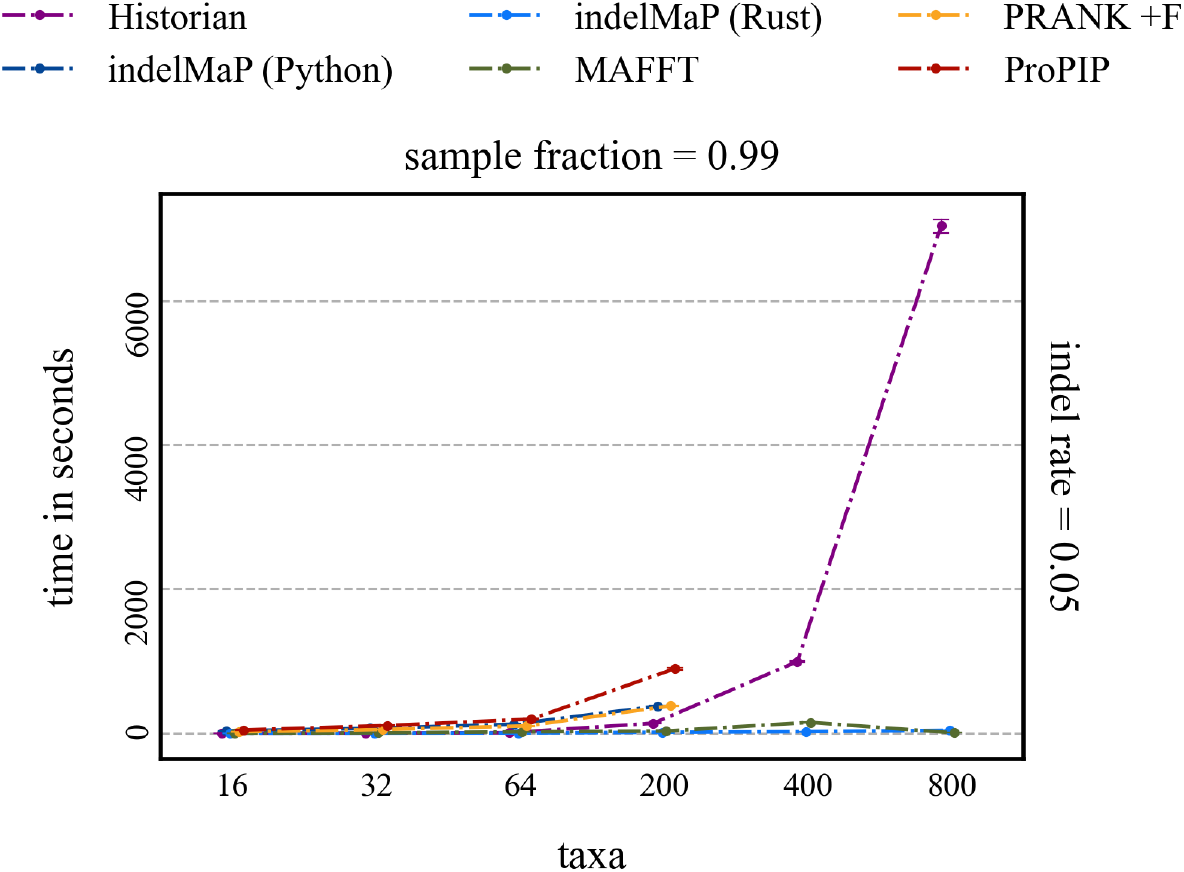
Computational time benchmark of MSA methods, for an extended data set with 400 and 800 taxa trees.

**Figure SM23:**
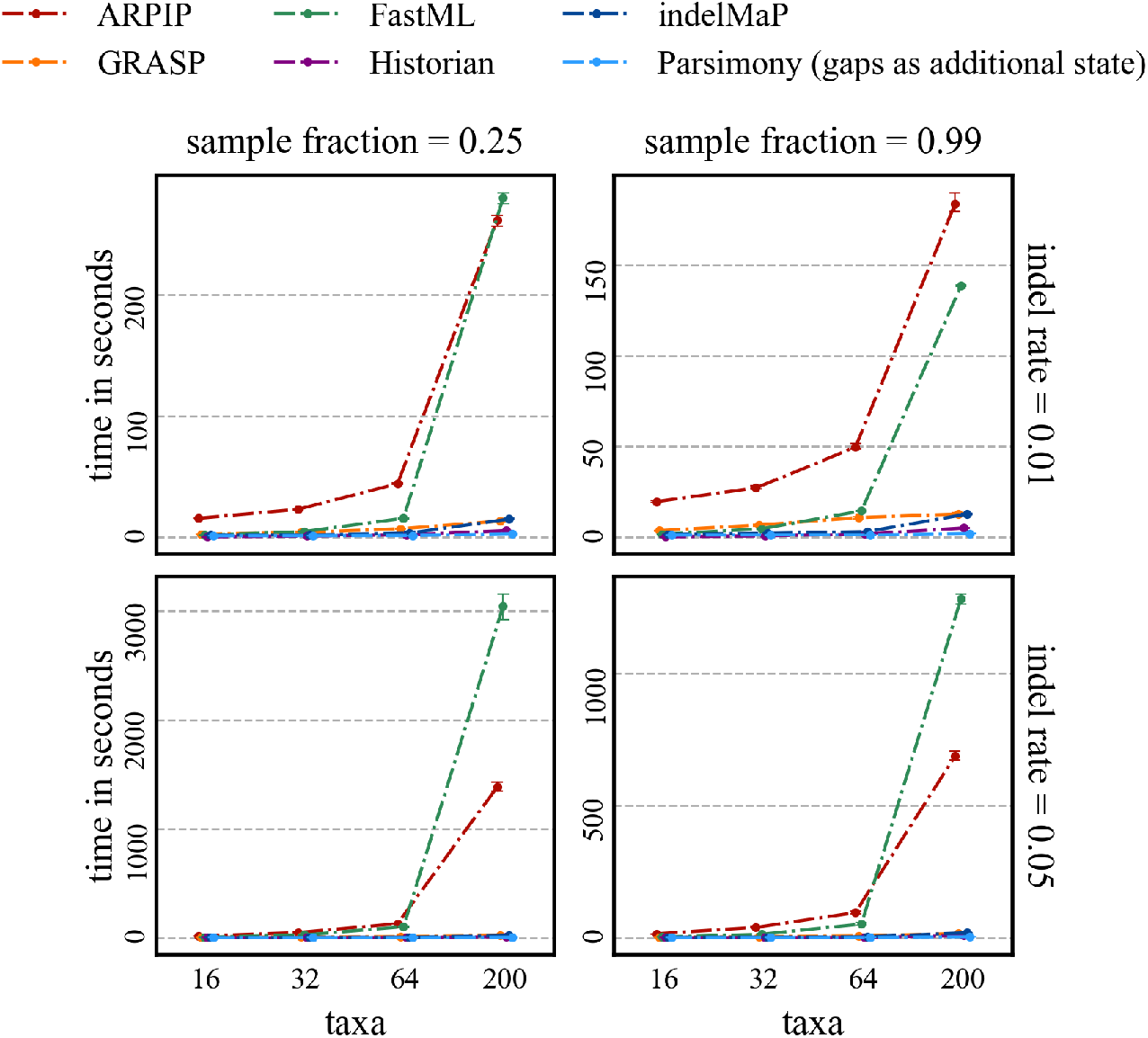
Computational time benchmark of ASR methods, under various parameter combinations with tree height 0.8.

**Figure SM24:**
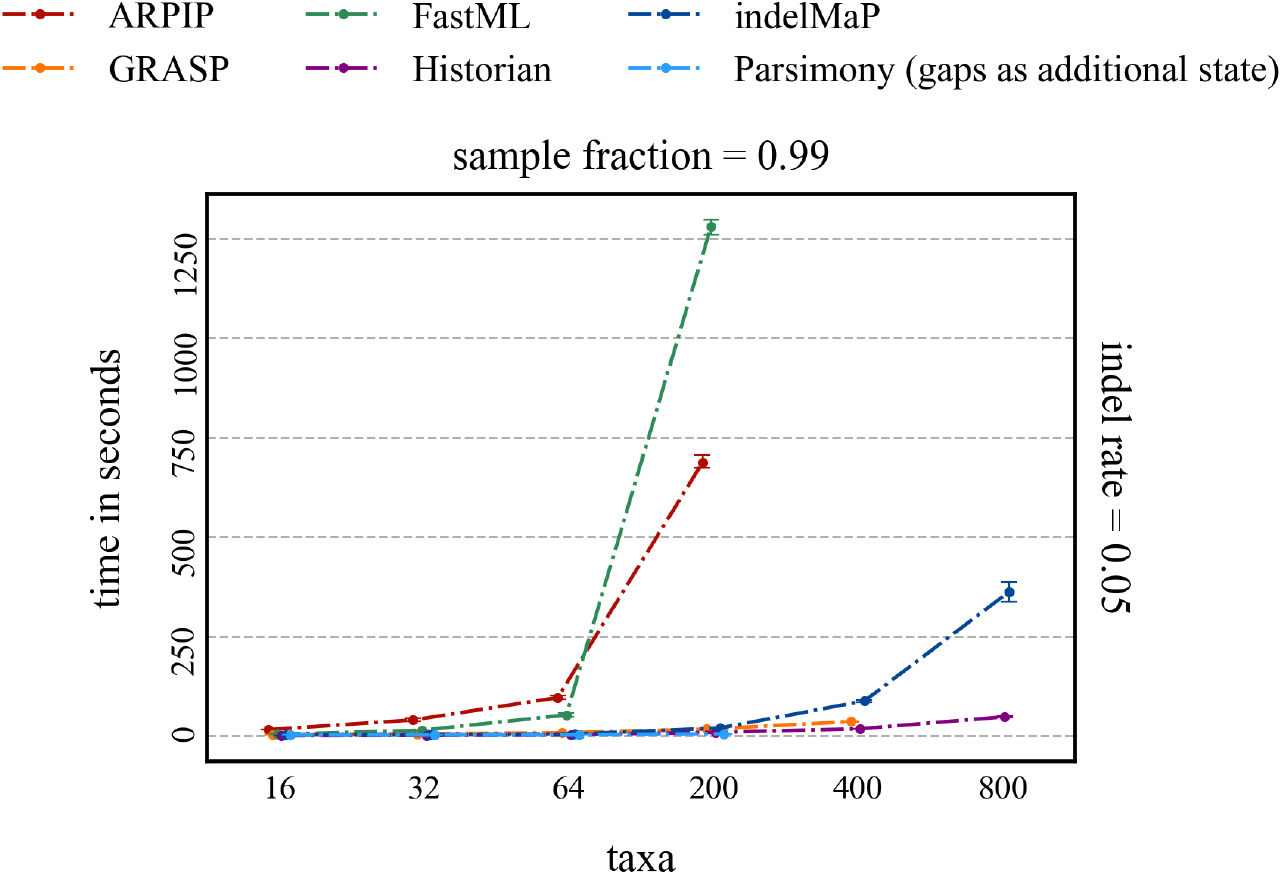
Computational time benchmark of ASR methods, for an extended data set with 400 and 800 taxa trees.

### A.8 Correlation between log-likelihood under PIP and indelMaP and Parsimony with gaps as an additional state for tree height 1.2

**Table SM1:** Pearson correlation between the log-likelihood score under PIP (log LK), indelMaP and the conventional parsimony score treating the gap character as an additional state (MP). All parameter combinations are for tree height 0.8.

|  | 16 taxa |  |  | 32 taxa |  |  | 64 taxa |  |  |  |
| --- | --- | --- | --- | --- | --- | --- | --- | --- | --- | --- |
|  | log LK | indelMaP | MP | log LK | indelMaP | MP | log LK | indelMaP | MP |  |
| log LK | - | -0.925 | -0.922 | - | -0.997 | -0.991 | - | -0.894 | -0.711 | 0.25 |
| indelMaP | -0.986 | - | 0.978 | -0.998 | - | 0.992 | -0.968 | - | 0.944 |  |
| MP | -0.975 | 0.982 | - | -0.99 | 0.993 | - | -0.817 | 0.927 | - |  |
| log LK | - | -0.948 | -0.92 | - | -0.896 | -0.863 | - | -0.988 | -0.971 | 0.99 |
| indelMaP | -0.983 | - | 0.993 | -0.969 | - | 0.992 | -0.987 | - | 0.985 |  |
| MP | -0.968 | 0.993 | - | -0.942 | 0.99 | - | -0.978 | 0.992 | - |  |

